# A single cell multi-omic analysis identifies molecular and gene-regulatory mechanisms dysregulated in the developing Down syndrome neocortex

**DOI:** 10.1101/2025.06.30.662136

**Authors:** Celine K. Vuong, Alexis Weber, Patrick Seong, Nana Matoba, Beck Shafie, Angelo Salinda, Pan Zhang, Yu-Jen Chen, Susanne Nichterwitz, Shahab Younesi, Le Qi, Michael J. Gandal, Daniel H. Geschwind, Jason L. Stein, Luis de la Torre-Ubieta

## Abstract

Down syndrome is the most common genetic cause of intellectual disability, presenting with cognitive, learning, memory, and language deficits. The cellular and molecular mechanisms driving this disorder remain unclear, limited by a lack of systematic studies in the developing human brain. Here, we leveraged single-nucleus multi-omics to profile the mid-gestation neocortex in a cohort of 26 donors. We observed a reduction in neural progenitors and corticothalamic neurons and concomitant increase of intratelencephalic neurons, accompanied by accelerated time to neuronal specification. We uncovered widespread changes in gene expression, chromatin accessibility and cell interaction networks impacting neurogenesis, specification and maturation and gene-regulatory networks directing these processes, including those downstream of transcription factors encoded in chromosome 21. Finally, we identified cell-specific molecular pathways shared with other neurodevelopmental disorders as well as heritability enrichment of GWAS signals in altered chromatin. Together, our data revealed a cascade of molecular dysregulation outlining the earliest steps in Down syndrome, providing a foundation for future therapeutic targets.

## Introduction

Down syndrome (DS) is the most common genetic cause of intellectual disability (ID), affecting ∼1 in 700 live births, and is caused by trisomy of human chromosome 21 (Ts21 of HSA21) ^1^. Individuals with DS show deficits in learning, memory and attention, delayed language and motor development, and atypical sensory processing ^1,2^. A subset of individuals with DS also have autism spectrum disorder (ASD, 10-15%), attention-deficit-hyperactivity disorder (ADHD, 20-44%) and/or epilepsy (8%), and virtually all are diagnosed with early-onset Alzheimer’s disease (AD) ^1,3,4^. Deficits in cognitive development appear as early as 6 months of age, and brain structural changes including reduced whole brain volume and reduced gyrification and sulcal depth in the temporal and frontal lobes, areas important for cognition, are already apparent during the second and third trimesters of development ^5–8^. The early emergence of cognitive defects coupled with reported changes in cortical structure suggest that corticogenesis, which peaks during mid-gestation ^9^, is likely disrupted in DS.

Previous studies investigating the impact of Ts21 on neurogenesis and specification have primarily employed model systems and revealed numerous impairments, but at times have produced conflicting results ^10^. These discrepancies likely reflect limitations in modeling the protracted length and complexity of neurogenesis, which occurs over the course of ∼3 months in vivo ^9,11^, species-specific differences in development ^9,12–14^ and incomplete construct validity of animal models ^15–18^. Furthermore, corticogenesis is governed by precise temporal and molecular processes that are still being refined in human in vitro model systems ^16,19,20^, altogether highlighting the need for human in vivo studies.

Transcriptome profiling of postmortem brains has advanced our understanding of molecular mechanisms driving DS, identifying altered programs in interneurons, microglia and oligodendrocytes ^21–23^. However, such studies have mainly focused on postnatal to adult periods, leaving a prominent gap in our understanding of the earliest steps leading to DS. The complexity of developing neocortex comprised of dozens of cell types with dynamic transition states and distinct genetic programs ^11,24–27^, including those regulating homotypic and heterotypic cell interactions that further modulate developmental decisions ^28,29^, necessitates single-cell resolution approaches. Transcription factors (TFs) and chromatin modifiers act on cis-regulatory elements (CREs) to direct gene-regulatory networks operating during corticogenesis ^24,26,30,31^, a process likely disrupted in DS ^32,33^. However, due in part to limited access to these critical tissues, paired transcriptomic and chromatin profiling of the developing Ts21 neocortex at single-cell resolution has not been systematically conducted.

To address this gap in connecting the gene regulatory processes dysregulated in DS to changes in neurodevelopment, we undertook single-nucleus, multi-omic profiling of gene expression and chromatin accessibility in developing Ts21 neocortex. We uncovered changes in cell composition resulting from an acceleration of the neurogenic timeline and altered excitatory neuron specification as a consequence of increased activity of the HSA21-encoded TF, BACH1. We leveraged paired multi-omic data to define gene expression programs and regulatory networks dysregulated in Ts21 and identify convergent risk genes and molecular pathways shared between DS and other neurodevelopmental disorders. Altogether, our work reveals neurodevelopmental changes occurring as early as mid-gestation, defines gene-regulatory mechanisms altered in Ts21, and provides an in vivo resource for future studies of DS neuropathology and gene regulation.

## Results

### A single-nucleus multi-omic analysis of the Ts21 neocortex at mid-gestation reveals widespread molecular dysregulation

To understand how Ts21 impacts cortical neurodevelopmental programs, we performed paired single-nucleus, multi-omic RNA sequencing and assay for transposase-accessible chromatin (ATAC) with sequencing (snMultiome) on a cohort of 13 control and 13 Ts21 neocortical samples from independent donors spanning the second trimester from gestational weeks (GW) 13 to 23 (**Figure 1A** and **Supplementary Table S1A**). We confirmed Ts21 and control karyotypes, using B-allele frequency (BAF) estimation (**Figure S1**). Control and Ts21 sample sets had similar mean ages and comparable distributions of sex, sample source, area and ancestry (**Figure 1B,C** and **Figure S2A,B**). After stringent QC, we retained 113,801 high quality nuclei (**Figure S2C**), performed cross-donor integration and clustered cells across all timepoints (see Methods). We annotated cell types by enrichment analysis to previously published datasets of developing neocortex and by expression of canonical marker genes (**Figure 1D,E** and **Figure S2D,E**; **Supplementary Table S1B**) ^25,26^, demonstrating high overlap of gene expression signatures across independent datasets.

**Figure 1.**
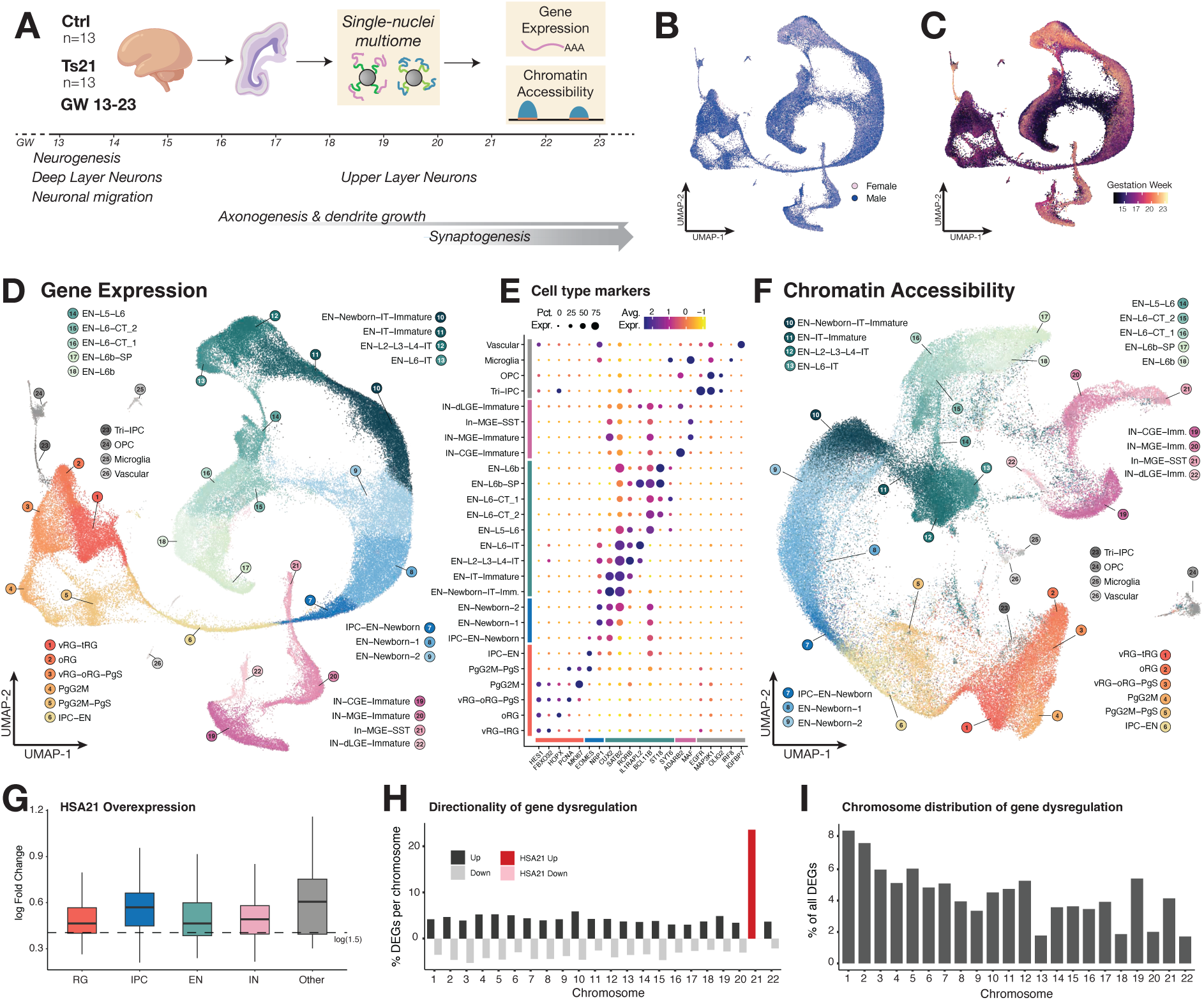
A multi-omic map of Down syndrome developing neocortex. (**A**) Schematic of the experimental design. Control and Ts21 samples from GW13-23 were dissociated into single nuclei and processed to obtain paired gene expression and chromatin accessibility. Major histogenetic processes occurring during profiled ages are listed along with their peak timing. (**B**) RNA-based UMAP showing distribution of nuclei from male (blue) and female (pink) samples. (**C**) RNA-based UMAP showing distribution of nuclei colored by gestation week. (**D**) RNA-based UMAP of cell types in this dataset. Clusters are colored by major cell type – Radial glia, orange; Newborn neurons, blue; Upper layer excitatory neurons, dark green; Deep layer excitatory neurons, light green; Inhibitory neurons, purple; Other cell types, grey. (**E**) Dotplot showing the expression of canonical marker genes for each major cell type. Dot size denotes percent cells in each cluster expressing the gene while color bar denotes average normalized expression. Individual color bars denote major cell types as in (D). (**F**) ATAC-based UMAP showing clustering by chromatin accessibility. Nuclei are colored by RNA-based annotation as in (D). **G**) Median HSA21 DEG overexpression in major cell classes. Dotted line denotes a 1.5 fold change. (**H**) DEGs as a percent of all genes encoded on each chromosome. Consistent with Ts21, HSA21 DEGs are only upregulated. (**I**) DEGs from each chromosome as a percent of all DEGs approximately follows chromosome size.

The resulting single-nucleus multi-omic atlas encompassed 26 clusters spanning all major cell types of the developing neocortex. These included six progenitor clusters (c1-c6) with radial glia (RG), intermediate progenitor cells (IPC) and their cycling states giving rise to three clusters of newborn neurons (c7-c9) transitioning into either corticothalamic (CT) and subplate (SP) deep layer neurons (c14-c18) or intratelencephalic (IT) neurons (c10-c13) at different stages of maturity (**Figure 1D**). We also captured four clusters of medial or caudal ganglionic eminence-derived (MGE or CGE) interneurons (IN) (c19-22) and the recently defined multivalent (capable of producing astrocytes, oligodendrocytes or interneurons) Tri-IPC progenitor ^26^ (c23), as well as oligodendrocyte precursor cells (OPCs, c24), microglia (c25) and vascular/endothelial cells (c26). Clustering was primarily defined by cell type identity, rather than donor, sex, or other covariates (**Figure 1B-D** and **S2B**) and we observed the expected change in cell type composition across developmental time - with RG, deep layer SP and CT neurons more abundant at early GWs while samples from later stages contained relatively more upper layer maturing IT neurons and INs (**Figure 1C,D**) ^9,11^. Chromatin accessibility profiles have been shown to be cell type-specific and change dynamically during neurogenesis similar to changes in transcriptional profiles ^24,26,30,31^. Indeed, we clustered nuclei using only their ATAC profiles and found that chromatin accessibility closely reflected transcriptomic changes (**Figure 1D,F** and **S2F,G**), with separation of the major cell types, including deep versus upper layer excitatory neurons (ENs) and MGE versus CGE INs. Finally, as expected, accessible peaks were enriched in functional gene-regulatory loci, including promoters and enhancers, rather than in heterochromatin (**Figure S2H).**

We aggregated differential expression or differential chromatin accessibility signatures across cell types and compared their distribution across chromosomes to systematically characterize gene expression and gene-regulatory differences in Ts21. As expected of the triplication, we observed that HSA21-encoded genes were exclusively upregulated at a 3:2 ratio (1.5x) (**Figure 1G; Supplementary Table S2**) consistent with previous brain gene expression studies ^21–23^. Only 25% of all HSA21-encoded genes were upregulated, pinpointing the most direct genetic drivers of the disorder (**Figure 1H)**. However, HSA21-encoded genes only comprised ∼5% of all differentially expressed genes (DEGs), indicating that Ts21 caused widespread transcriptional changes (**Figure 1I**). Similar to DEGs, nearly all detected peaks within HSA21 (∼80%) were more accessible, consistent with an extra copy of the chromosome, yet reflected only ∼40% of all differentially accessible (DA) peaks (**Figure S2F,G**). These results validate prior observations using bulk or single-cell gene expression in the adult brain ^21,23^ and extend them to the developing neocortex, along with newly-generated matching chromatin architecture changes. Dysregulated genes included many transcriptional and epigenetic regulators, such as those known to interact with the BAF, REST and PRC2 families of developmental chromatin remodelers (*BRWD1*, *DYRK1A*, *HMGN1*; **Supplementary Table S2**) ^33–35^, and genes involved in asymmetric division and synaptogenesis. Given the central role of gene regulation in the ordered generation and development of the neocortex, we hypothesized that gene dysregulation in Ts21 could lead to major changes in neurogenesis and neuronal development of the DS brain and sought to characterize this in depth.

### Altered neurogenesis and excitatory neuron specification in Ts21 neocortex

As histology and brain imaging studies have reported gross cytoarchitectural abnormalities in Ts21 ^6,36,37^, we leveraged our dataset to ascertain cell composition in detail. Control and Ts21 samples clustered together and were composed of the same major cell classes and cell types (**Figure 2A**). To assess changes in composition, we first quantified the proportion of each major cell class (RG, EN, IN and Other non-neuronal cells) over all timepoints. A small increase in INs was observed, consistent with previous work ^21^, but since their progenitors were not profiled we focused on the EN lineage. We observed a significant reduction in RGs suggesting altered neurogenesis but no differences in overall EN proportion (**Figure S3A**), prompting further investigation at the cluster level. We found that most progenitor clusters, except oRGs, were significantly decreased in Ts21 (**Figure 2B**), consistent with class-level changes. Among neurons, deep layer CT ENs (EN-L6-CT_2, EN-L6b-SP) were significantly decreased, while IT ENs (EN-Newborn-IT-Immature, EN-IT-Immature, EN-L2-L3-L4-IT, EN-L6-IT) showed a concomitant increase. To understand cell composition dynamics during corticogenesis, we quantified cells across early (GW13-15), mid (GW16-18) and late (GW19-23) periods of mid-gestation (**Figure 2C**). Ts21 apical progenitors (vRG-tRG, vRG-oRG-PgS) showed a reduction over developmental time. In contrast, basal (oRG) and intermediate progenitors (IPC-EN and IPC-EN-Newborn), which normally expand at later stages of corticogenesis and primarily give rise to upper layer neurons ^11^, increased at early/mid timepoints and decreased precociously in Ts21. The neurons that arise from these progenitors followed a similar pattern - deep layer CTs (EN-L6-CT_2) were decreased early in Ts21 while upper layer ITs (EN-L2-L3-L4-IT) showed a steady increase during mid and late periods. We also observed a significant increase in Tri-IPCs, a progenitor cell type that normally appears at later gestational ages ^26,38^. These changes are consistent with an acceleration of the neurogenic timeline, which stereotypically first produces deep layer neurons followed by upper layer neurons.

**Figure 2.**
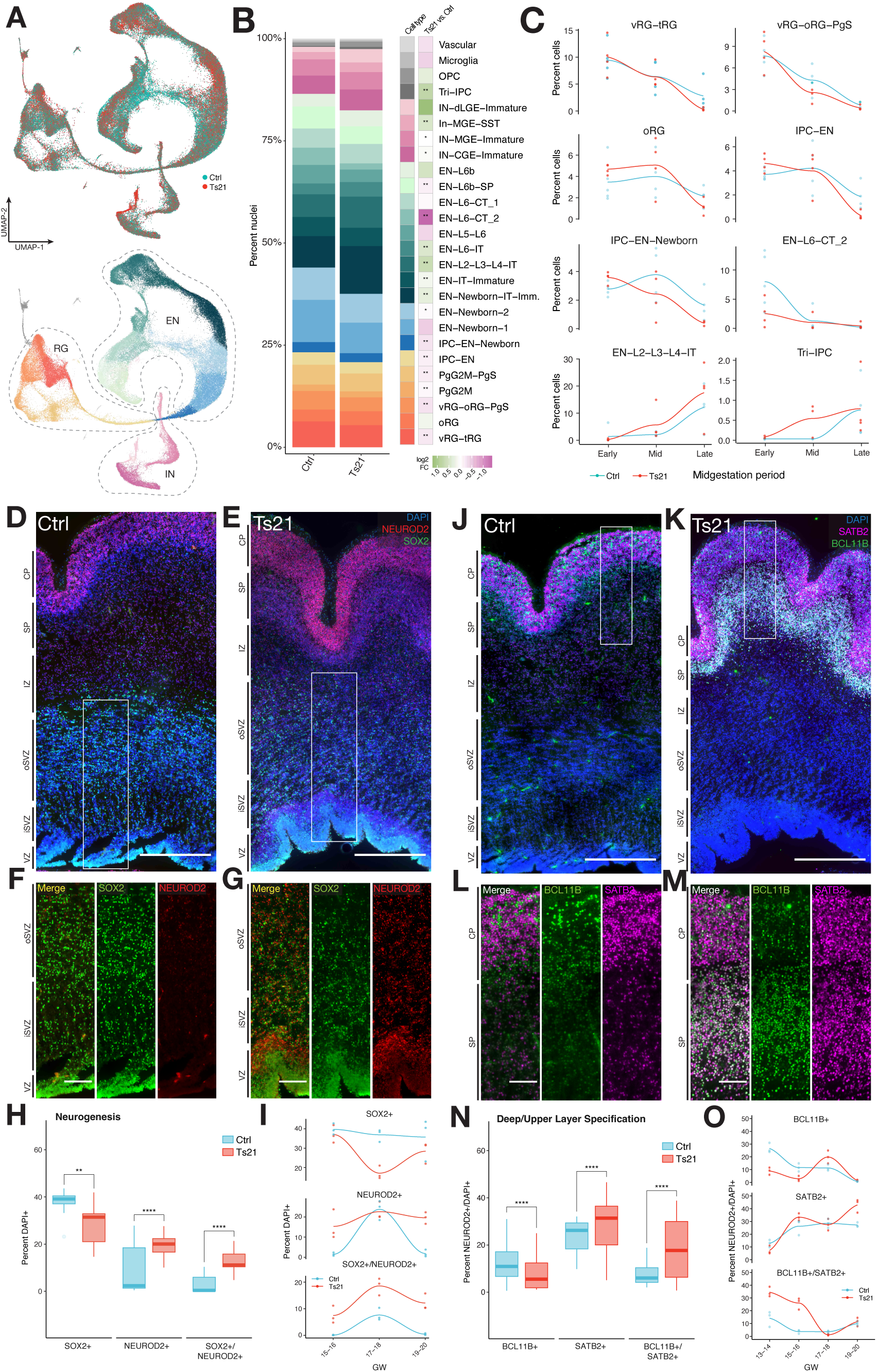
Altered cell composition and excitatory neuron specification in DS. (**A**) Top, RNA-based UMAP showing cell type distribution of control and Ts21 nuclei. Bottom, UMAP denoting major cell classes. (**B**) Cluster-level cell type composition over all time points shows significant decrease in multiple RG cell types, cycling progenitors, and deep layer CT EN, and significant increase in upper layer IT ENs, INs and Tri-IPC in Ts21 compared to control. FDR-corrected *P<0.01, **P<0.001. (**C**) Select cell type composition changes over time. Donor samples were binned into Early (GW13-15), Mid (GW16-18) and Late (GW19-23) periods. Each point is the average percent composition of samples from each donor. (**D**, **E**) Immunostaining of NEUROD2 (red) and SOX2 (green) in neocortex from age-matched GW15 control and GW16 Ts21 samples. DAPI, blue. Scalebar, 500um. (**F**, **G**) Insets as denoted in (D, E) encompassing VZ to oSVZ. NEUROD2/SOX2 double positive cells, yellow. Scalebar, 150um. (**H**) Percent of cells across all time points and cortical layers that are NEUROD2+ only, SOX2+ only or double positive. Overall, SOX2+ cells are significantly decreased while NEUROD2+ or double positive cells are significantly increased in Ts21. (**I**) Percent NEUROD2+, SOX2+ or double positive cells at each time point. (**J**, **K**) Immunostaining of SATB2 (magenta) and BCL11B (green) in neocortex from age-matched GW15 control and GW16 Ts21 samples. DAPI, blue. Scalebar, 500um. (**L**, **M**) Insets as denoted in (J, K) encompassing SP and CP. SATB2/BCL11B double positive cells, white. Scalebar, 100um. (**N**) Percent of excitatory neurons across all time points and cortical layers that are BCL11B+ only, SATB2+ only or double positive. BCL11B+ cells are significantly decreased while SATB2+ or double positive cells are significantly increased in Ts21. (**O**) Percent SATB2+, BCL11B+ or double positive excitatory neurons at each time point. FDR-corrected **P<0.01, ***P<0.001, generalized linear models (H, N). VZ, ventricular zone; iSVZ, inner subventricular zone; oSVZ, outer subventricular zone; IZ, intermediate zone; SP, subplate; CP, cortical plate.

To validate our transcriptomic findings, we performed immunohistochemistry (IHC) in cortices at developmentally matched timepoints (GW13-21) across a set of 8 donors (n=4 control, n=4 Ts21). To overcome potential technical biases and the low throughput of manual quantification, we developed an automated pipeline to quantify samples at scale (**Figure S3B-E**). We first assessed whether the ratio of neural progenitors to neurons was altered in Ts21 using the respective progenitor and excitatory neuron markers SOX2 and NEUROD2 (**Figure 2D,E**). Across all timepoints, we found that the percent of SOX2+ cells in the proliferative layers (VZ, iSVZ, oSVZ; **Figure 2F,G**) was significantly decreased in Ts21, with a concomitant increase in both NEUROD2+ and SOX2+/NEUROD2+ double positive cells (generalized linear model, FDR-corrected P-value<0.05, **Figure 2H,I**), consistent with the reduction in multiple RG cell types detected by transcriptomic analysis. Next, we used the respective deep and upper layer markers BCL11B and SATB2 to assess EN specification (**Figure 2J,K**) within the neuronal layers (IZ, SP, CP; **Figure 2L,M**). We found a significant decrease in BCL11B+ and concomitant increase in both SATB2+ and SATB2+/BCL11B+ double positive neurons (FDR-corrected P-value<0.05, **Figure 2N**) over all timepoints, consistent with our observation that upper layer IT neurons are increased at the expense of deep layer neurons. Similar to transcriptomic changes, the reduction in BCL11B+ neurons in Ts21 occurred at earlier time points (GW13-14) while the increase in SATB2 occurred later (GW15-21; **Figure 2O**). We also observed a significant increase in BCL11B+/SATB2+ double positive cells at early time points (GW13-16), potentially indicating mis-specification of CT versus IT neurons in Ts21. We then compared the average expression of sets of canonical upper layer IT (*SATB2*, *CUX2*, *RORB*) and deep layer CT (*BCL11B*, *TLE4*, *ZFPM2*) TFs across all cell types to ascertain whether gene programs important for establishing and maintaining CT and IT identity ^39^ are mis-expressed. In Ts21, upper layer TFs were significantly upregulated across all deep layer CT and SP cell types (EN-L5-L6, EN-L6-CT_1/2, EN-L6b and EN-L6b-SP; linear mixed models, FDR-corrected P-value<0.05; **Figure S3F**), while deep layer TFs were increased in multiple IT cell types (EN-Newborn-IT-Immature, EN-IT-Immature, EN-L2-L3-L4-IT, EN-L6-IT; **Figure S3G**) and decreased in some deep layer cells (EN-L5-L6). Together our observations point to an acceleration of the neurogenic timeline in Ts21 leading to a reduction of the progenitor pool and early production of IT neurons at the expense of CT neurons, as well as mis-specification of IT and CT neurons.

### Heterochrony in the Ts21 neocortex leads to accelerated neurogenesis and neuronal maturation

To understand the dynamics of cell composition changes in the developing Ts21 neocortex, we inferred the differentiation trajectory of cells within the EN lineage (**Supplementary Table S3**) using Monocle3 ^40^. Using the earliest developmental time point of GW13 vRG-oRG-PgS cells as the root node, we ordered cells by their pseudotime along the EN lineage. As expected, RGs had the lowest pseudotime and highest transcriptional similarity to the root, with progressively increased pseudotime and transcriptional difference in IPCs and newborn neurons, followed by deep and upper layer neurons, which diverge into two separate lineages (**Figure 3A**).

**Figure 3.**
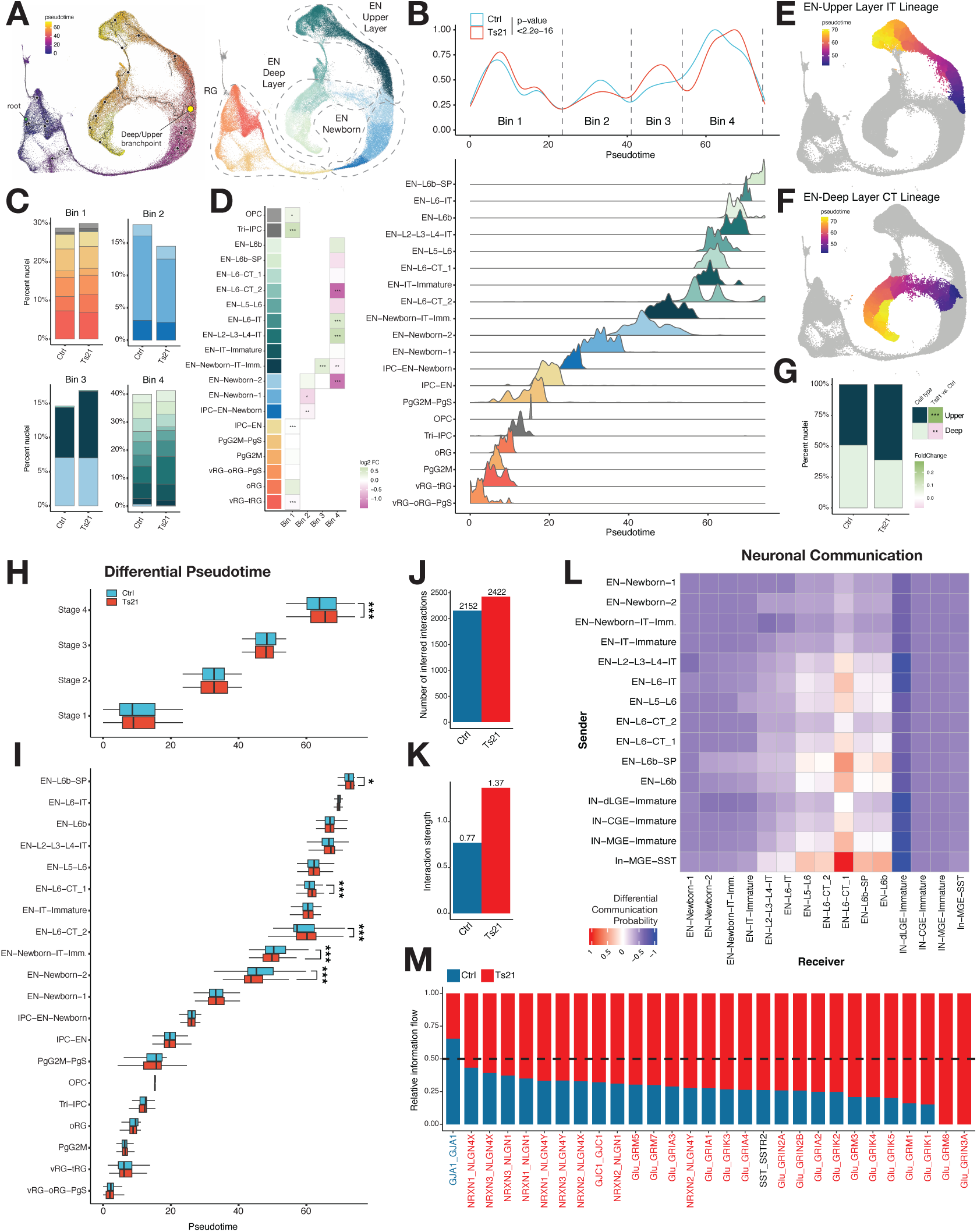
Accelerated neurogenesis and neuronal maturation in DS. (**A**) Left, RNA-based UMAP of pseudotime trajectory of the excitatory neuron lineage. The root (GW13 vRG-oRG-PgS) node is labeled in green, and the deep/upper layer neuron branchpoint is labeled in yellow. Right, UMAP showing major cell types. (**B**) Top, density plot of cells over pseudotime shows that the Ts21 distribution is significantly different compared to control. Kolmogorov-Smirnov test. Dotted lines denote pseudotime bins. Bottom, ridge plot of cell type distributions over pseudotime. (**C**, **D**) Differential cell type composition of each pseudotime bin in Ts21 versus control. Generalized mixed models. (**E**, **F**) Pseudotime UMAP of upper (E) and deep (F) layer EN lineages. (**G**) Deep versus upper layer cell composition in Ts21 compared to control. (**H**, **I**) Differential pseudotime of each pseudotime bin (H) or cell type (I). Deep layer EN-L6-CT_1/2 and EN-L6-SP show significantly increased pseudotime in Ts21, while EN-Newborn-IT-Imm. and EN-Newborn-2 have significantly decreased pseudotime. FDR-corrected *P<0.05, **P<0.01, ***P<0.001. (**J, K**) NeuronChat analysis. The number (J) and strength (K) of neuronal interactions is increased in Ts21. (**L**) Heatmap displaying differential communication probability between neurons. (**M**) Neuronal communication pathways significantly increased in Ts21 (red) include multiple NRXN-NLGN interactions and glutamatergic signaling. Pathways significantly decreased in Ts21 are in blue, non-significant pathways in black (FDR-corrected P<0.05).

Mapping cell densities along pseudotime, we observed that control cells accumulated into four bins: Bin 1 – RG, Bin 2 – IPCs and newborn neurons, Bin 3 – immature neurons, Bin 4 – maturing deep and upper layer neurons; (**Figure 3B**; **Supplementary Table S3**). While Ts21 cells followed an overall similar distribution, cell density was lower in Bin 2 and higher in Bin 3. In contrast, Bins 1 and 4 had similar density but showed a relative shift towards increased pseudotime in Ts21, altogether leading to significantly different pseudotime distribution between conditions (Kolmogorov-Smirnov test; P<2.2e16; **Figure 3B**). To understand differentiation dynamics, we quantified cell composition (**Figure 3C**), finding significant changes at all pseudotime bins (**Figure 3D**; FDR-corrected P<0.05, linear mixed models). In Bin 1, Ts21 vRG-tRG and IPC-EN were significantly decreased while Tri-IPC and OPC were increased, consistent with our previous observations (**Figure 2B**). Next, Ts21 cells showed a significant decrease of IPC-EN-Newborn and EN-Newborn-1 in Bin 2 and an increase of EN-Newborn-IT-Immature in Bin 3, reflecting an acceleration through Bin 2, where cells do not yet express neuronal subtype markers, and accumulation in Bin 3, where expression of *SATB2* and *NKAIN2* indicate commitment to the IT lineage. Finally, Ts21 neurons showed a significant decrease in CT (EN-L6-CT_2) and increase in IT (EN-L2-L3-L4-IT, EN-L6-IT) neurons in Bin 4 as previously observed (**Figure 2B**). Separating EN cells at the branchpoint where CT and IT lineages diverge (**Figure 3 A,E,F**), we found that while CT and IT neurons were evenly distributed in control, this ratio was significantly altered in favor of IT neurons in Ts21 (39% deep layer, 61% upper layer; FDR-corrected P-value<0.05, linear mixed models; **Figure 3G**).

The cell composition differences at pseudotime Bins 2 and 3 point to changes in the rate of neuronal commitment and differentiation. To test this we calculated differential pseudotime, finding that Ts21 cells in Bin 4 had significantly increased pseudotime (**Figure 3H**, FDR-corrected P<0.001, linear mixed models). Similarly, at the cell type level deep layer CT neurons (EN-L6-CT_1 and EN-L6-CT_2, P<0.001; EN-L6b-SP, P<0.05) showed significantly increased pseudotime with upper layer IT neurons (EN-L2-L3-L4-IT) also showing a trending increase. Interestingly, pseudotime of EN-Newborn-2 and EN-Newborn-IT-Immature cells was significantly decreased (P<0.001). This is consistent with an acceleration in neurogenesis, as increased numbers of newly committed neurons would initially show a less differentiated transcriptional profile. Reduced pseudotime, in combination with the observed accumulation of Ts21 cells in Bin 3 (**Figure 3B**) and an increase in EN-Newborn-IT-Immature cells at the same stage (**Figure 3C,D**) suggests that the transition from progenitor to newborn neuron is accelerated in Ts21, leading to a relative increase in newly committed neurons.

To understand how expression of HSA21 genes might contribute to differentiation dynamics, we assessed their expression over pseudotime (**Figure S4**). We found that HSA21 gene expression is spread over all stages of pseudotime, consistent with changes in both progenitors and neurons. Notably, a number of genes located at the pseudotime transition between progenitor and neuron include HSA21-encoded TFs *BACH1* and *GABPA* whose peak expression precede that of neuronal subtype-specific TFs such as *CUX2*, *SATB2* and *BCL11B*, suggesting an additional mechanism affecting CT versus IT neuron specification. An acceleration of the neurogenic timeline would result in more mature neurons in the Ts21 neocortex, as demonstrated by their increased pseudotime. To assess neuronal maturation using an orthogonal approach, we performed neuron-neuron communication analysis using NeuronChat ^41^. We found that both the number and strength of interactions were increased in Ts21 (**Figure 3J,K**; **Supplementary Table S4A,B**), and that communication probability within ENs as well as between ENs and INs (**Supplementary Table S4C**) was increased. In particular, ligand-target interactions altered in Ts21 originated from both deep and upper layer neurons targeting EN-L6-CT_1 and from MGE-SST targeting deep layer ENs (**Figure 3L**). Significantly increased pathways included many well-established synaptic interactions such as NRXN-NLGN ^42^ and glutamatergic signaling (Glu-GRIA, Glu-GRM) ^43,44^ (**Figure 3M**, **Supplementary Table S4D**; FDR-corrected P<0.05). At our profiled timepoints synaptogenesis is at its earliest stages and highly dynamic ^45,46^, thus the observed increase in neuron-neuron interactions suggests that Ts21 neurons are more mature, consistent with an earlier transition to neurogenesis rather than altered circuit connectivity. Together, these changes in trajectory and neuronal signaling demonstrate an acceleration of neurogenesis and neuronal maturity consistent with proposed heterochrony in DS ^10^ as well as altered specification of IT and CT neurons in Ts21.

### Molecular mechanisms controlling metabolism and neurogenesis are altered in Ts21

We next sought to understand the gene expression changes underlying altered neurogenesis in Ts21. Using a rigorous hierarchical, multi-sample mixed model (see Methods) ^47^ we identified 3,464 cell type-specific DEGs, of which 2,065 (59%) were significantly upregulated and 1,399 (41%) were significantly downregulated (FDR-corrected P-value<0.05, **Figure 4A**; **Supplementary Table S2**). Of the 278 detected HSA21 genes, 145 (52%) were significantly upregulated, none were downregulated, and HSA21 genes comprised ∼25% of the DEGs identified in each cell type (**Figure S5A**), consistent with previous findings ^21,23^. We observed significant gene dysregulation in most cells, but ENs, both IT and CT, had the largest number of DEGs while relatively fewer changes were observed in progenitors, INs and non-neuronal cells (OPC, Microglia, Vascular), the latter likely due to the smaller proportion of those cells in the neocortex. To identify molecular pathways affected during the major stages of neurodevelopment in Ts21, we first pooled cell type-specific DEGs by class (RG, IPC, EN, and IN) and performed gene ontology analysis.

**Figure 4.**
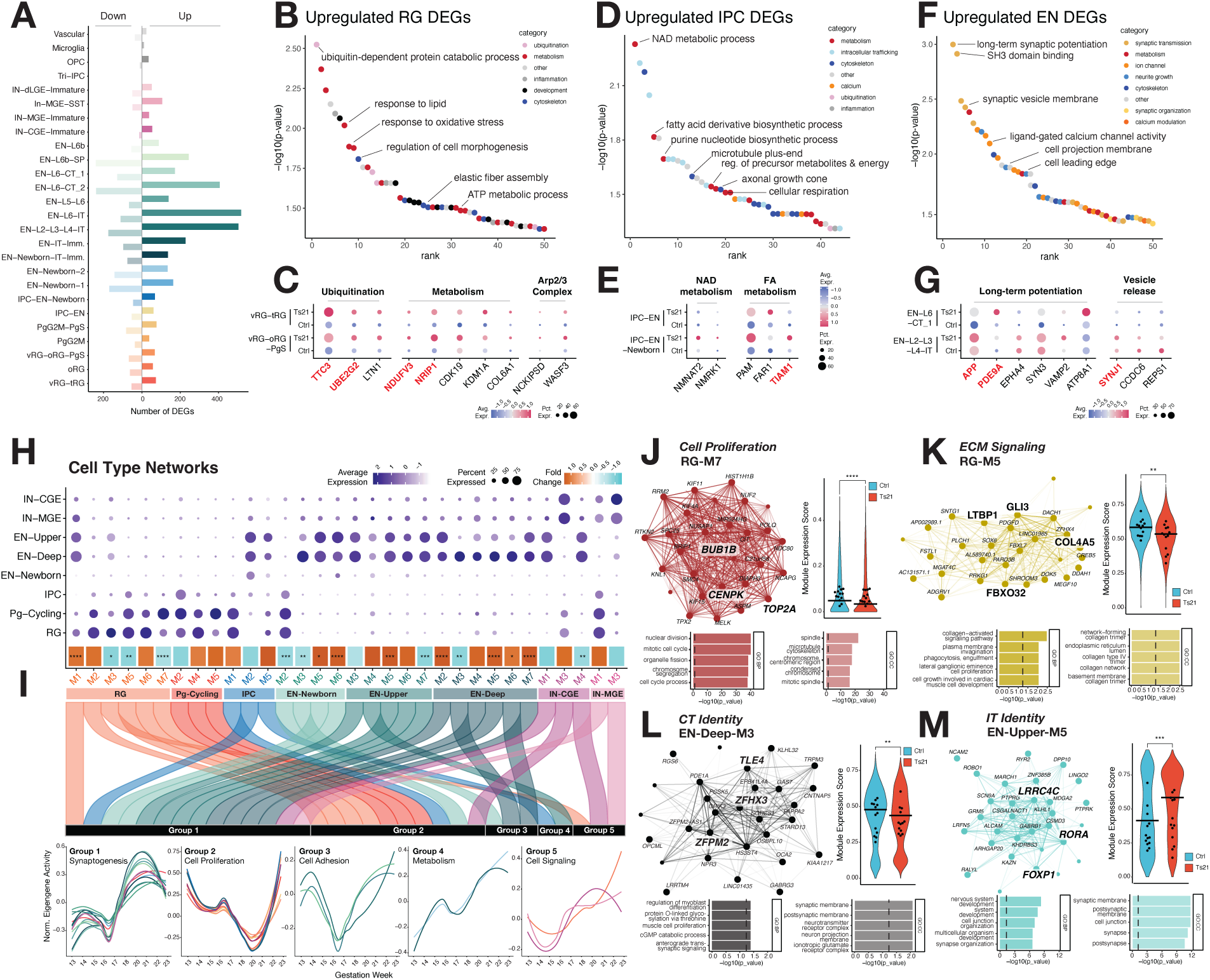
Pro-neurogenic and altered neuronal specification programs in Ts21. (**A**) Number of differentially expressed genes per cell type. Excitatory neurons show the most dysregulation, followed by newborn neurons, neural progenitors and inhibitory neurons. (**B-G**) Gene ontology of upregulated cell type-specific DEGs grouped by cell class. (**B, D, F**) Top 50 GO terms ranked by P-value. Terms are colored by category, with specific GO terms highlighted. (**C, E, G**) Expression of specific genes associated with processes upregulated in RGs, IPCs and ENs, respectively. HSA21-encoded genes are highlighted in red. (**H**) Cell type-specific gene co-expression modules. Top, dotplot showing expression of module eigengenes across major cell types. Average fold change of module expression in control vs Ts21 is shown below dotplot. **I**) Alluvial plot grouping cell type-specific modules into 5 major categories. Normalized eigengene expression of grouped modules over developmental time is shown below. (**J-M**) Select module network plots showing hub genes (top left), average eigengene expression score in control vs Ts21 (top right), and top 5 biological process (BP) and cell compartment (CC) gene ontology terms enriched in each module (bottom). Vertical dashed line indicates FDR-corrected P-value<0.05. (**J**) RG-M7 involved in cell cycle and proliferation is significantly decreased in Ts21. (**K**) RG-M5 involved in proliferative ECM-mediated signaling is downregulated.( **L, M**) EN-Deep-M3 and EN-Upper-M5 regulating CT and IT identity are respectively down- and up-regulated in Ts21. FDR-corrected *P<0.05, **P<0.01, ***P<0.001.

Ts21 RGs exhibited upregulation of genes enriched in processes related to ubiquitination, metabolism and cytoskeletal dynamics, among others (**Figure 4B**, FDR-corrected P-value<0.05). Changes in ubiquitination and metabolism were driven in part by several HSA21 genes. The E3 ligase *TTC3*, which targets phosphorylated Akt leading to reduced proliferation ^48^, was upregulated along with other components of the ubiquitin proteasomal pathway (*UBE2G2*, *LTN1*; **Figure 4C**). Two metabolic HSA21-encoded genes - *NDUFV3*, a component of oxidative phosphorylation (OXPHOS), and *NRIP1*, part of the glycolytic pathway - were also significantly upregulated, likely contributing to metabolic imbalance and increase in genes involved in mitochondrial function (*CDK19*, *KDM1A/LSD1*, *COL6A1*; **Figure 4C**) ^49–51^. While mitochondrial dysfunction has been documented in Ts21 along with mis-expression of both OXPHOS and glycolytic genes ^10,52,53^, we found more mitochondrial function genes upregulated in RGs overall, consistent with accelerated neurogenesis as neural progenitors switch from glycolysis to OXPHOS during differentiation ^54,55^. Genes regulating cytoskeletal dynamics were also altered in Ts21 RGs, including two associated with the Arp2/3 complex (*NCKIPSD*, *WASF3*; **Figure 4C**), a key component in promoting neurogenic asymmetric division ^56^.

Similarly to RGs, IPCs displayed pro-neurogenic gene expression changes in metabolism, including upregulation of processes supporting OXPHOS such as NAD biogenesis (*NMNAT2*, *NMRK1*) and fatty acid metabolism (*PAM*, *FAR1*; **Figure 4D,E**) ^54,57,58^. Changes in cytoskeletal dynamics in IPCs also promote neuronal differentiation, driven in part by upregulation of HSA21-encoded *TIAM1*, a GTPase activator important for neuronal polarity and axonogenesis (**Figure 4E**) ^59^. Gene expression changes in ENs reflected accelerated neurogenesis in Ts21, with increased synaptogenesis and maturation (**Figure 4F**) driven in part by HSA21-encoded genes such as *APP* and *PDE9A* which promote development of long-term synaptic potentiation ^60,61^ along with other non-HSA21 genes (*EPHA4*, *SYN3*, *VAMP2*, *ATP8A1*; **Figure 4G**). Similarly, synaptic vesicle release and recycling genes are upregulated, including HSA21-encoded *SYNJ1* and non-HSA21 genes such as *CCDC6* and *REPS1* (**Figure 4G**). Upregulated IN DEGs were primarily related to metabolic processes similar to RGs, IPCs and ENs, and included OXPHOS and FA metabolism genes (*NDUFV3*, *FAR2*; **Figure S5B,C**).

While we initially observed few to no pathways enriched in downregulated genes, increasing the number of pooled DEGs using a lower cutoff (FDR-corrected P-value<0.1) revealed enrichment of processes disrupted in progenitor maintenance, neuronal connectivity and RNA metabolism (**Figure S5D**). In RGs, downregulated genes were enriched in lysosomal autophagy (*SERAC1*, *ATG14*, *STX17*), a process impaired in neurodegeneration ^62^ but which has been recently shown to also influence asymmetric division ^63^; concordantly, genes involved in fatty acid elongation, a necessary step in lysosomal formation ^64^, were downregulated in oRGs (*ACOT7*, *ELOVL6*, *ELOVL2*; **Figure S5E**). In IPCs and ENs, downregulated genes were enriched in processes related to axo-dendritic polarization and axonogenesis (*LRP8*, *CAV3*, *EZR*, *SEMA3A*, *SHTN1*; **Figure S5F,G**) ^65–68^ consistent with reports of polarization deficits in Ts21 neurons ^22^. RNA metabolism was also dysregulated in ENs, with reduced expression of mRNA degradation (*PARN*, *CNOT1*, *CNOT2*) ^69^ and stability ^70^ (*SYNCRIP*) factors in EN-L2-L3-L4-IT (**Figure S5G**). While changes in splicing have been studied in Ts21 tissues including in the brain ^21,22^, dysregulation of other aspects of RNA metabolism have not been previously described. We found genes downregulated in ENs were enriched in chromatin remodeling, specifically in EN-L6-IT neurons where expression of genes such as *BAZ1A*, a member of the ISWI complex, and *NIPBL* were reduced (**Supplementary Table S2**). Finally, IN DEGs were enriched in dendritic and post-synaptic development (**Figure S5D**) consistent with reported dysregulated inhibitory function in DS mouse models ^71^ and reduced incoming neuronal communication probability (**Figure 3L**). Overall, Ts21 DEGs regulate the balance between proliferation and neurogenesis and affect key aspects of neuronal differentiation important for proper connectivity and function.

### Gene co-expression network analysis reveals dysregulated neuronal specification and maturation programs in Ts21

We next performed high-dimensional weighted gene co-expression network analysis (hdWGCNA), a system-level approach that uses unsupervised learning to identify dysregulated pathways based on co-expression structure and is inherently more powered than DEG analysis ^72^ (see Methods). We built gene co-expression networks for each of the 8 major cell types – RG, Pg-Cycling, IPC, EN-Newborn, EN-Deep Layer, EN-Upper Layer, IN-MGE and IN-CGE – and assigned genes to modules based on shared patterns of co-variation across samples (**Figure 4H**; **Supplementary Table S5 A-H**). After filtering (see Methods), we retained 32 modules related to major corticogenesis processes including proliferation (RG-M2/M5/M7), neurogenesis (RG-M6, EN-Newborn-M2/M6), EN specification (EN-Newborn-M3/M5, EN-Upper-M5, EN-Deep-M3/M5/M6), neuronal maturation (EN-Newborn-M5, EN-Upper-M6, EN-Deep-M4) and synaptic development (EN-Newborn-M5/M6, EN-Upper-M7, EN-Deep-M2/M7) (**Supplementary Table S5**). While module expression was primarily cell type-specific, as expected we observed overlapping expression between related cell types such as RG and Pg-Cycling, Deep and Upper layer neurons, and IPCs with progenitors and their neuronal progeny (**Figure 4H**, **Figure S5H**). To identify gene programs altered in Ts21, we calculated the gene activity of each module and compared the resulting scores across conditions, finding that 16/32 expressed modules (50%) were significantly differentially expressed (**Figure 4I**, **Supplementary Table S5I**). We then classified modules across networks into five major groups based on their expression pattern over time and gene ontology – Group 1, Synaptogenesis; Group 2, Cell Proliferation; Group 3, Cell Adhesion; Group 4, Metabolism; Group 5, Cell Signaling (**Figure 4I** and **Figure S5I**).

Consistent with our DEG analyses, we found that modules promoting proliferation and symmetric division (Group 2) were all significantly downregulated, including cell division RG-M7 (hub genes *BUB1B*, *CENPK*, *TOP2A*; **Figure 4J**) ^73–75^ and Newborn-M2 (hub genes *ZEB1*, *GAB1*; **Figure S5J**) which suppresses asymmetric division ^76,77^. Interestingly, pro-proliferative extracellular matrix (ECM)-mediated signaling in RGs was also dysregulated as evidenced by decreased activity of RG-M5 (**Figure 4K**) containing RG-specific TFs (hub genes *GLI3*, *DACH1*, *ZFHX4*) and ECM components mediating TGFb signaling (hub genes *LTBP1* and *COL4A5*) ^29,78,79^. In contrast, the largest module group consisted of programs related to synaptogenesis (Group 1), where 6/7 differential modules were upregulated in Ts21 (Newborn-M5/M6, EN-Upper-M5 and EN-Deep-M2/M5/M6) (**Figure 4H,I**). Of these, 5 modules regulating pre-and post-synaptic signaling and organization were expressed in both deep and upper layer neurons (Newborn-M5/M6, EN-Upper-M5, EN-Deep-M2/M7; **Figure 4M**, **Figure S5K**), consistent with accelerated neurogenesis and DEG analyses.

Finally, we uncovered changes in molecular programs regulating the establishment and refinement of EN identity. In Group 3, we found two modules associated with the establishment of L6-CT neuron identity, selectively expressed in deep layer neurons and containing key CT TFs as hub genes ^39^ (Newborn-M3 – *ZFPM2*, Deep-M3 – *TLE4*, *ZFPM2*; **Figure S5L**, **Figure 4L**), were significantly downregulated. Conversely, two modules with IT-associated hub genes ^80–82^ (Newborn-M5 – *SATB2*, EN-Upper-M5 – *RORA*, *LRRC4C*, *FOXP1*; **Figure S5K**, **Figure 4M**) were expressed in both CT and IT neurons and upregulated in Ts21. These gene program changes are consistent with our observation that the IT/CT neuron ratio is increased in Ts21 and that key IT and CT transcription factors are mis-expressed (**Figure S3F,G**), resulting in the mixed-identity state we observe with increased SATB2+/BCL11B+ neurons (**Figure 2J-O**). In addition to promoting CT identity, ZFPM2 and TLE4 suppress a related corticofugal neuron subtype, SCPN, and loss of these TFs results in mixed corticofugal identity and mistargeting of L6 axons ^83^. Concordantly, we found that two modules with SP and SCPN hub genes expressed across all deep layer cell types ^25,84–86^ (EN-Deep-M5 – *PCP4*, *CELF4*; EN-Deep-M6 – *ST18*, *LMO3*, *CDH18*; **Figure S5M**) were significantly upregulated. EN-Deep-M5 and -M6 were enriched in genes associated with post-synaptic signaling ^44,87^ (*IGSF2*, *GABRB1/2*, *RYR3*) and axon pathfinding ^88,89^ (*SLIT3*, *ARHGAP4*4), suggesting that dysregulation of corticofugal programs could affect L6 neuronal connectivity and circuit function in Ts21.

### Altered vascular endothelial-RG interactions promote neurogenesis in Ts21

Disrupted ECM-mediated signaling in Ts21 RGs suggested that inter-cellular mechanisms could be implicated in neurogenic changes. Within the neurogenic niche, extrinsic signaling from RGs, neurons, vascular endothelial cells and ECM interactions regulate neuronal progenitor maintenance and differentiation ^28,29,90,91^. To explore whether inter-cellular signaling is disrupted in Ts21, we performed cell-cell communication analysis using CellChat ^92^ (see Methods). We identified 67 total ligand-receptor pathways (**Supplementary Table S6**) of which 50/67 (82%) showed significantly differential ligand-receptor pathway activity, with 9 pathways downregulated and 41 pathways upregulated (P-value<0.05, **Figure 5A**). Synaptic signaling pathways in ENs and INs (Glutamate, GABA-A and GABA-B; **Figure 5A**, **Figure S6**) were significantly upregulated in Ts21, similar to our observations using NeuronChat (**Figure 3M**) and consistent with accelerated neurogenesis and neuronal maturation. In addition, we observed significantly increased interactions between neurons in pathways important for neurite outgrowth and patterning ^93,94^ (SEMA6, CDH, CNTN, EPHB, Netrin, CADM, SLIT and SLITRK; **Figure 5A**) consistent with accelerated maturation.

**Figure 5.**
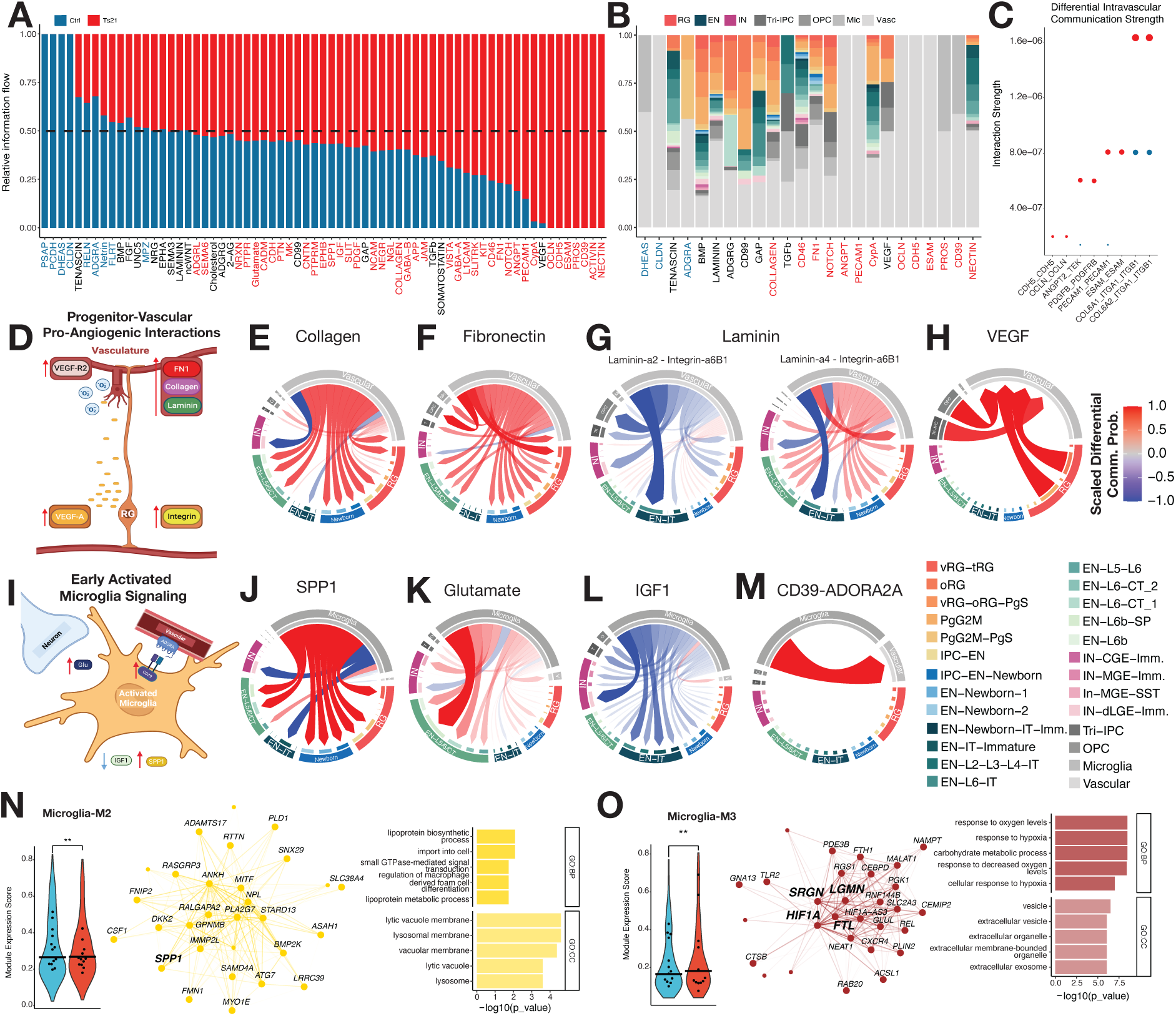
Pro-angiogenic and inflammatory microglial signaling are increased in Ts21. (**A**) CellChat RankNet plot of 67 pathways in the merged network: 9 downregulated (blue), 41 upregulated (red) in Ts21 (FDR-corrected P-value<0.05). (**B**) Cell type RankNet plot: 24 pathways enriched in vascular and microglia (>25%, P=0.0018); 3 in control, 13 in Ts21. (**C**) Intravascular interaction strengths: vascular ligand-receptor (LR) pairs show increased or exclusive signaling in Ts21 (red) compared to control (blue). (**D**) Overview of RG-Vascular pro-angiogenic interactions: VEGFA (left, orange) secretion by RGs, recruits endothelial tip cell ingression through vascular VEGFR2 (left, pink). Angiogenesis and RG asymmetric division are accelerated by increased RG integrin (right, yellow) binding basal end-feet to vascular cell ECM (Fibronectin/FN1, red; Laminin, green; Collagen, purple). (**E-H**) Differential chord plots of RG-Vascular ECM interactions. Cell sender is at the origin of the arrow, cell receiver is at the end of the arrow. Arrows are colored by feature scaling of the differential communication probability between pathway LR pairs (blue -1 lower in Ts21, red +1 higher in Ts21). (**E,F**) Increased collagen and fibronectin/FN1 signaling in Ts21. (**G**) Proliferative laminin α2-integrin signaling is reduced (left) while pro-neurogenic laminin α4-integrin is increased. (**H**) VEGFA-VEFGR signaling between RGs and vascular cells is increased in Ts21. (**I**) Overview of pro-inflammatory microglia signaling: hallmarks of early activated microglia including glutamate (Glu, purple) release to neurons (blue), increased SPP1 and decreased IGF1. (**J-M**) Differential chord plots of microglia interactions. (**J,K**) Increased SPP1 signaling from microglia to multiple cell types (J) and glutamate signaling from microglia to neurons (K), indicative of a pro-inflammatory state, are increased. (**L**) IGF1 signaling from microglia, a marker of an anti-inflammatory state, is decreased. (**M**) Increased pro-angiogenic adenosine signaling via microglia-expressed CD39 and vascular-expressed ADORA2A. (**N**,**O**) Microglia co-expression network modules showing hub genes (left), average eigengene expression score in control vs Ts21 (center), and top 5 biological process (BP) and cell compartment (CC) gene ontology terms enriched in module (right). Modules M2 and M3 containing microglia activation markers as hub genes (bolded) and regulating phagocytic processes are significantly increased in Ts21. FDR-corrected **P<0.01. [Figure created with BioRender]

We noted that in 16/50 (32%) of significantly different pathways, vascular cells and microglia made up 25% or more of the ligand-receptor interactions (**Figure 5B**) despite comprising a minority of the cells in the dataset, suggesting vascular signaling was disrupted in the Ts21 neocortex (**Figure 5B** and **Figure S6**). Neurogenesis is closely tied to vascular development, as the signaling between RGs, endothelial cells and perivascular cells promotes angiogenesis and vascular ingression required to oxygenate the developing brain ^90,91,95,96^. We found that tight junction proteins (OCLN, PECAM1, ESAM, and CDH5; **Figure 5C**) important for blood vessel growth and integrity ^97,98^ – and other intra-vascular pro-angiogenic signaling interactions including ANGPT2-TEK ^99,100^, Jagged1-Notch ^101^ and PDGFB-PDGFRB ^102^ (**Figure 5C**) were upregulated in Ts21.

We also observed significantly increased signaling between vascular cell ECM, including collagen (COL4A1, COL6A1, COL6A2, COL1A1) and fibronectin with RG integrin-β1 (**Figure 5D-F)**, which promote RG end-feet adhesion to perineural vascular plexus at the pial surface required for asymmetric RG division ^96,103–108^. In addition, we observed pro-neurogenic laminin signaling between vascular cells and RGs ^109^ - laminin-ɑ2-integrin signaling, which promotes RG attachment to the ventricular surface and proliferative maintenance ^110^, was downregulated while laminin-ɑ4-integrin signaling, required for RG attachment to the pial surface necessary for neuronal migration ^111^, was upregulated (**Figure 5D,G**). In addition to altered ECM signaling, Ts21 RGs showed increased VEGF signaling to vascular cells (**Figure 5D,H**), a well-established pro-angiogenic signaling pathway promoting vessel ingression towards the VZ ^95^. Altogether intra-vascular and RG-vascular signaling in Ts21 promote a pro-neurogenic and pro-angiogenic environment consistent with increased OXPHOS gene programs, reduced proliferation and overall accelerated neurogenesis.

### Pro-inflammatory microglia signaling is increased in Ts21

Increased microglial activation in post-mortem studies and models of DS has been linked to early-onset neurodegeneration as well as excessive synaptic pruning ^112–116^, but it has not been described in developing brain. We observed significant increases in microglia-enriched inflammatory signaling pathways consistent with an activated microglial state in Ts21 including PTN (pleiotrophin) ^117^, MK (midkine) and SPP1 (osteopontin) ^118^, among others (**Figure 5A,B**). We found that Ts21 microglia increased SPP1 signaling (**Figure 5I,J**), which promotes phagocytic states ^119^, as well as glutamate release (**Figure 5I,K**), an indicator of an activated state ^120,121^. Conversely, IGF1 secretion (**Figure 5I,L**) indicating an anti-inflammatory state ^122,123^ was downregulated. Ts21 microglia also showed increased signaling of the NTPDase CD39, indicative of an activated state, to vascular ADORA2A (**Figure 5I,M**) which could promote vascular proliferation and angiogenesis ^124,125^. To assess microglial states using an orthogonal method, we generated a microglia gene co-expression network using hdWGCNA and compared the expression of modules across conditions (**Supplementary Table S5J,K**). Of the 3 expressed modules, 2 were significantly upregulated (FDR-corrected P-value<0.05), contained pro-inflammatory hub genes (Microglia-M2 - *SPP1*, *GPNMB*, *DKK2*; Microglia-M3 -*FTL*, *SRGN*, *LGMN* and *HIF1-a*) and were enriched in phagocytic and activation processes (**Figure 5N,O**). Increased *SPP1*, *SRGN* and *HIF1-a* expression has been associated with activated or phagocytic microglia ^126,127^, while elevated levels of *FTL* and *LGMN* have been observed in neurodegeneration ^128,129^, suggesting that microglial activation may begin as early as prenatal stages in Ts21.

### Widespread chromatin dysregulation in Ts21

To characterize gene-regulatory dynamics in Ts21, we integrated paired gene expression and chromatin accessibility modalities using the weighted nearest-neighbor ^130^ (WNN, see Methods) method and performed joint clustering, identifying 27 WNN cell types (**Figure 6A**) similar to the 26 identified using only gene expression (**Figure 1D**). Chromatin accessibility closely reflected gene expression in a cell type-specific manner. All cell types as defined by gene expression were discernible in similar cell clusters generated using only chromatin accessibility (**Figure S7A**), and we observed strong concordance between gene expression, gene activity (chromatin accessibility at the TSS) and motif activity of known cell type-specific TFs (**Figure S7B**).

**Figure 6.**
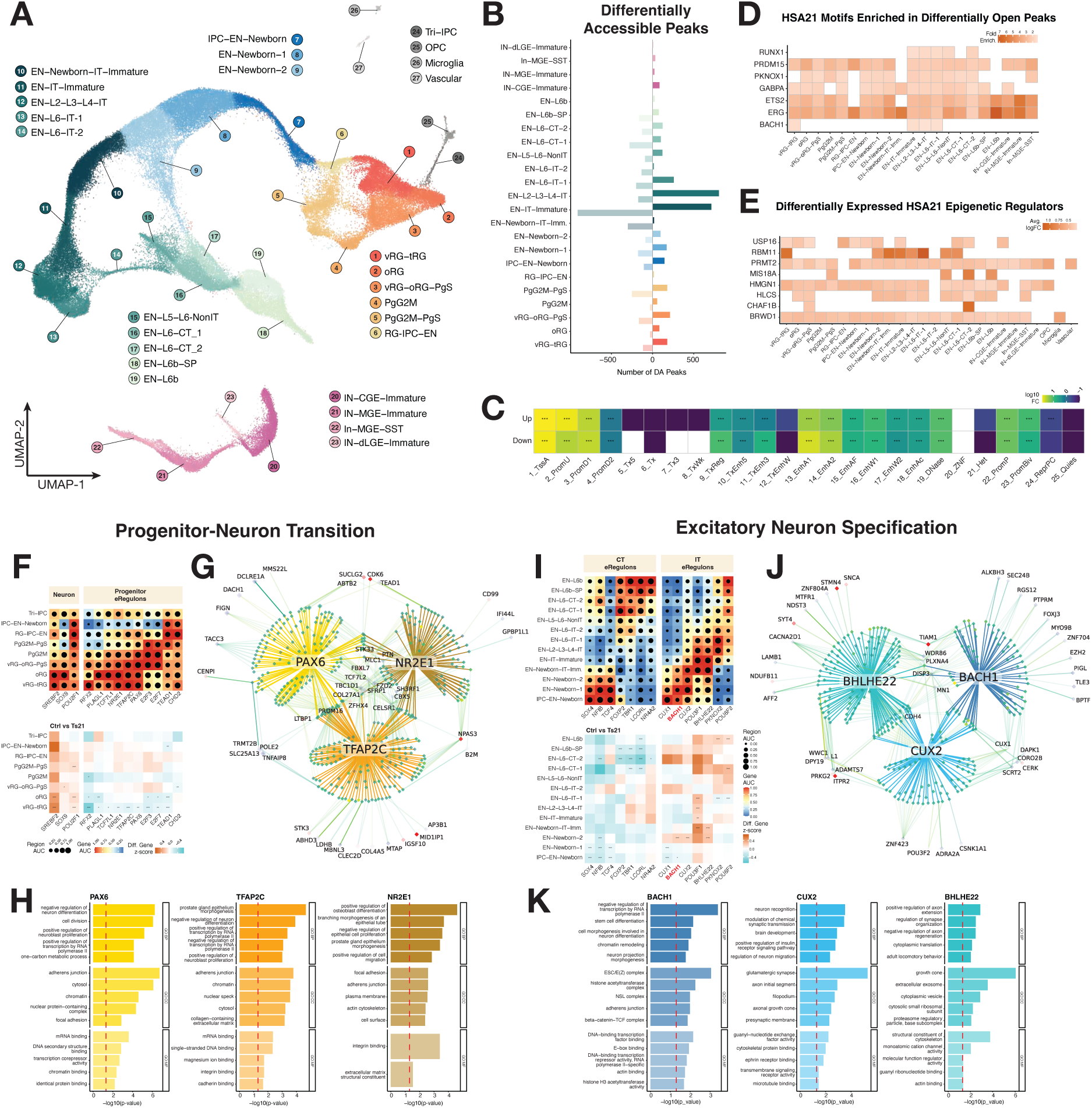
Widespread chromatin dysregulation and upregulation of pro-neurogenic and pro-IT GRNs in Ts21. (**A**) WNN-based UMAP of cell types jointly clustered by RNA and ATAC. Clusters are colored by major cell type – Radial glia, orange; Newborn neurons, blue; Upper layer excitatory neurons, dark green; Deep layer excitatory neurons, light green; Inhibitory neurons, purple; Other cell types, grey. (**B**) Cell type-specific DA peaks. Excitatory upper layer neurons (EN-IT-Immature, EN-L2-L3-L4-IT) have the largest number of DA peaks. (**C**) GREAT enrichment of annotated gene-regulatory regions in DA peaks. Color bar shows fold change, FDR-corrected ***P-value<0.001. (**D**) Heatmap of cell type-specific motif enrichment of HSA21-encoded TFs in differentially open peaks. Only significant (FDR-corrected P<0.05) enrichments are shown. Color bar denotes fold enrichment. (**E**) Heatmap of cell type-specific HSA21-encoded DEGs known to function as epigenetic regulators. Only significant changes are shown. Color bar denotes average log fold change. (**F-H**) Select eRegulons illustrating pro-neurogenic gene-regulatory programs active in Ts21. (**F**) Top, heatmap dotplot of pro-neurogenic or pro-progenitor eRegulon activity (AUC, area under the curve) by target gene expression (color bar) and chromatin accessibility (dot size) in RGs. Bottom, heatmap of differential eRegulon gene activity. Color bar denotes differential activity z-score. Two-sided t-test, FDR-corrected *P<0.05, **P<0.01, ***P<0.001. (**G**) PAX6, TFAP2C and NR2E1 pro-proliferation eRegulons. Primary edges from TFs (yellow, orange, brown) denote significant TF-to-region connections and inner green diamonds denote chromatin regions. Secondary edges from regions denote significant region-to-gene (R2G) connections, with darker colors and thicker edges indicating a stronger R2G score. Outer diamonds denote eRegulon genes targeted by each region; red color denotes increased gene expression in Ts21. (**H**) Top 5 biological process (BP), cell compartment (CC) and molecular function (MF) gene ontology terms enriched in selected eRegulon target genes. Red dotted line denotes FDR-corrected P-value<0.05. (**I-K**) Select eRegulons illustrating pro-IT neuron gene regulator programs increased in Ts21. (**I**) Top, heatmap dotplot of pro-CT and pro-IT eRegulon activity. Bottom, heatmap of differential activity of CT and IT eRegulons. HSA21-encoded TF BACH1 is colored in red. (**J**) CUX2, BHLHE22 and BACH1 pro-IT eRegulons. (**K**) Gene ontology terms enriched in pro-IT eRegulons.

We next identified cell type-specific DA peaks, finding that nearly all cell types except for Tri-IPCs, OPC, microglia and vascular cells exhibited significant changes in chromatin accessibility consistent with DEG results (**Figure 6B** and **Figure S7C**; **Supplementary Table S7A**) (**Figure 4A**). Similar to gene expression, chromatin accessibility changes were highest in ENs and particularly upper layer neurons (EN-IT-Immature, EN-L2-L3-L4-IT). Ts21 DA peaks were enriched in promoters and enhancers, including those with poised or bivalent epigenetic marks ^131,132^, and under-enriched in heterochromatin, indicating a role in gene regulation (**Figure 6C**, GREAT method ^133^ binomial test, FDR-corrected P-value<0.05, -log10 fold-change>0). We then asked which TF motifs were enriched in DA peaks (**Figure S7D**; **Supplementary Table S7B)** and found that those of the largest TF families - C2H2 zinc finger, homeodomain and BHLH, with established roles in neurogenesis and enriched in dynamic GZ/CP chromatin changes ^24^ - were the most abundant, suggesting altered neurogenesis gene-regulatory programs in Ts21. Focusing on HSA21-encoded TF motifs enriched in Ts21>control DA peaks, we found that 7/9 known motifs were significantly enriched (**Figure 6D**; **Supplementary Table S7B**). While some were over-represented in all cell types (PRDM15, PKNOX1, GABPA, ETS2, ERG), two were specifically enriched in excitatory neurons (RUNX1, BACH1) (**Figure 6D**). RUNX1 enrichment in active EN chromatin is consistent with its known roles in promoting neuronal differentiation and peak expression in the most mature cells (**Figure S4**). While BACH1 has been shown to function in microglia and astrocyte differentiation in the developing brain and as a contributor to neurodegeneration in the adult brain, its role in neurogenesis has not been well explored. Widespread chromatin changes in the Ts21 neocortex could also be driven by HSA21-encoded epigenetic regulators, which have been shown to modulate genome-wide transcriptional changes ^33,34,134,135^. Indeed we observed that multiple chromatin modifiers including *BRWD1*, *HMGN1* and *PRMT2* were significantly upregulated across many cell types (**Figure 6E**). Together these suggested that transcriptional dysregulation in Ts21 could also be driven by altered chromatin accessibility leading to differential availability of CREs.

### Increased activity of gene-regulatory networks promoting neurogenesis and IT specification in Ts21

We next sought to conduct a systematic analysis of gene-regulatory networks (GRNs) impacted in Ts21. We applied SCENIC+ ^136^ to link TFs to their putative enhancers and target genes, identifying 358 enhancer-driven regulons (eRegulons; 196 activators, 162 repressors; **Supplementary Table S8A**) with cell type-specific gene and region (chromatin) activity. Fidelity of the GRNs was demonstrated by the capture of well-established cell type-specific TF activity in the expected cell types (**Figure S7E)**. We identified 5 eRegulons centered on HSA21-encoded TFs (GABPA, BACH1, OLIG2, ETS2 and ERG; **Figure S7E**). OLIG2, ETS2 and ERG showed highly specific expression in OPC, microglia and vascular cell types, respectively, while GABPA and BACH1 were active in RG, newborn neurons and INs. Focusing on activators, we identified 90 eRegulons with significantly differential activity in Ts21 in at least one cell type (see Methods, two-sided t-test, FDR-corrected P-value<0.05, **Supplementary Table S8B**; **Figure S8**). Below, we highlight several sets of eRegulons that illustrate changes occurring in GRNs controlling the transition from progenitor to neuron, and the specification of CT versus IT excitatory neurons in the developing Ts21 cortex.

Across multiple RG cell types (oRG, vRG-tRG), eRegulons promoting stemness over differentiation were significantly downregulated (**Figure 6F**). These included the RG markers PAX6 and TFAP2C, which promote RG identity ^137,138^ and cell division ^30,139^ (**Figure 6G**), transcriptional regulators controlling cell cycle such as NR2E1/TLX ^140,141^, E2F3 ^142^ and E2F7 ^143^ and the chromatin remodeler CHD2, which suppresses the canonical neuronal repressor REST/Nrsf ^144,145^ (**Figure 6F**). Interestingly, we noted significantly reduced RG activity of 3 eRegulons involved in pro-proliferative signaling occurring at the primary cilium ^146^ – TCF7L1, which promotes RG stemness via Wnt-beta-Catenin ^147^; TEAD1, a component of the Hippo pathway ^148,149^; and RFX2, a coordinator of ciliary gene expression ^150^ (**Figure S9A**). Disrupted cilia signaling is consistent with recent work demonstrating altered ciliogenesis caused by aberrant expression of the HSA21 gene *PCNT* ^151,152^, which we find significantly upregulated in RGs and ENs (**Supplementary Table S2**). Finally we found reduced vRG-tRG and oRG activity of PLAGL1 (**Figure 6F**, **Figure S9A**), a regulator of the apical to basal RG transition ^153^ driving ECM-mediated signaling ^154^, consistent with observed dysregulation of ECM expression programs (**Figure 4K**) and accelerated neurogenesis (**Figure 2B**, **Figure 3B-D**). Upregulated GRNs included SOX9, which regulates expression of ECM components required for oRG expansion and later gliogenesis ^78,155^, POU2F1/OCT-1, whose role in cortex is unknown but which controls the timing and specification of retinal progenitors ^156^ and SREBF2, enriched in target genes involved in cholesterol biosynthesis (**Figure 6F**, **Figure S9B**) consistent with its known role as a major regulator of this process ^157,158^. Interestingly, recent work has shown that a cholesterol gene expression program is associated with the transition from apical (vRG) to basal (oRG) radial glia^159^.

We next sought to define GRNs driving the cell specification phenotypes observed in the excitatory lineage. Previous studies have identified key TFs influencing the ordered progression of EN specification into first CT then IT neurons ^26,39,80,160–165^, including recent work identifying eRegulons acting at branch points along CT/IT lineages ^26^. From these studies we compiled 33 CT/IT eRegulons and found that 27/33 (81%) were active during mid-gestation, with 14/33 (42%) significantly dysregulated in Ts21 (**Figure 6I**). Overall, differential eRegulons reflected the shift in proportion of CT to IT neurons we observed using cell type composition analyses (**Figure 2**), with 7/7 CT eRegulons significantly decreased in newborn or CT neurons, and 6/7 IT eRegulons significantly increased in newborn, CT and IT neurons (**Figure 6I**). These included TFs known to function in deep layer CT specification, identity and migration downregulated in newborn neurons (SOX4, NFIB and TCF4 ^26,160,166^; IPC-EN-Newborn, EN-Newborn-1/2; **Figure 6I**, **Figure S9C**), maturing CT (FOXP2, TBR1 ^167,168^; EN-L6-CT_1/2; **Figure 6I**) and SP (NR4A2/NURR1 ^163^; Figure 6I). Conversely, eRegulons promoting upper layer IT identity were upregulated in newborn and maturing IT (CUX1, CUX2 ^164,169^ in EN-Newborn-2; POU3F1, BHLHE22 ^26,165,170^ in EN-Newborn-2, EN-Newborn-IT-Immature, EN-L2-L3-L4-IT, EN-L6-IT-1; Figure 6I) as well as mis-expressed in CT and non-IT (PKNOX2, POU6F2 ^26^; EN-L5-L6-NonIT, EN-L6-CT_2, EN-L6b; Figure 6I). Finally, we asked whether altered CT/IT specification involved GRNs driven by HSA21-encoded TFs. We observed that the BACH1 eRegulon active in newborn and immature neurons was significantly upregulated in EN-Newborn-2, one of the earliest clusters showing commitment to the IT lineage (Figure 3). Target genes of BACH1 included other TFs promoting IT identity such as *CUX2* and *BHLHE22*; consistent with this, *BACH1* peak expression precseded that of *CUX2* in pseudotime (**Figure S4**), suggesting that altered neuron specification is due in part to direct activity of this HSA21-encoded gene. Enrichment of BACH1 motifs was also observed in Ts21 DA peaks within ENs in the companion study ^171^, suggesting this dysregulation extends to postnatal periods.

### Convergence of neurodevelopmental pathways altered in Ts21 with rare and common genetic signals of neuropsychiatric disorders

As the most common cause of ID with a known genetic origin, Ts21 provides a model for understanding pathways shared across conditions with cognitive impairment. Moreover, given the co-occurence of DS with other neurodevelopmental or neuropsychiatric conditions, including ASD, ADHD and epilepsy, we sought to investigate whether genetic signals for ID or these conditions overlapped with transcriptional and/or chromatin alterations identified in Ts21. We first conducted enrichment analyses to test if rare-variant association signals from large-scale whole-exome or whole-genome sequencing studies across ASD ^172^, NDD ^172^, DD ^173^, and epilepsy ^174^ overlapped with cell-specific DEGs or modules altered in Ts21 (Figure 7A). For ASD, we also tested syndromic and non-syndromic variants from the curated SFARI database. We observed significant overlap of multiple cell-specific DEGs and modules across most disorders. Upregulated DEGs and gene programs enriched in disease gene sets were primarily involved in neuronal differentiation and synaptic function, reflecting accelerated neurogenesis and maturation in Ts21. As such, we focused on those signals downregulated in Ts21 to match the expected haploinsufficiency caused by *de novo* rare variants. Genes downregulated in EN-L6-CT-2 showed consistent enrichment across multiple disease gene sets (**Supplementary Table S9A**), including NDD (FDR P = 4.94e-05), DD (FDR P = 2.49e-04), ASD (FDR P = 3.75e-04), and syndromic ASD (FDR P = 2.12e-03). Similarly, those downregulated in EN-L6-IT were significantly enriched for NDD (FDR P = 2.03e-03) and DD (FDR P = 2.76e-04) risk genes. These findings implicate deep-layer excitatory neurons as a point of vulnerability across developmental disorders, motivating further examination of the intersecting gene sets (Figure 7B).

**Figure 7.**
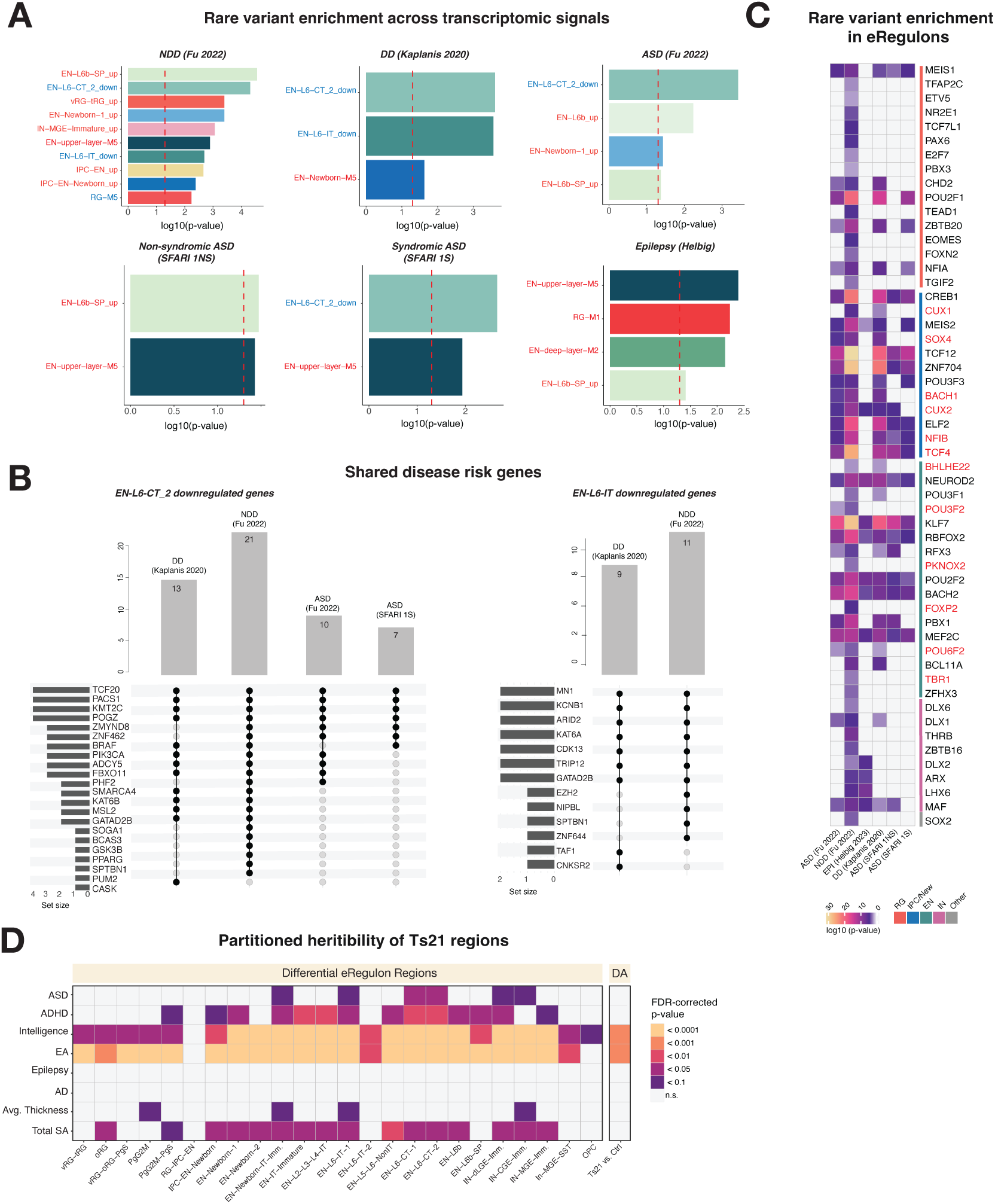
Convergent dysregulation of Ts21-altered neurodevelopmental pathways with rare and common genetic signals of neuropsychiatric disorders. (**A**) Rare variant enrichment across Ts21-dysregulated transcriptomic signatures. Enrichment of cell-specific upregulated (red) and downregulated (blue) cell type-specific DEGs and gene co-expression modules with NDD, DD, ASD, and epilepsy-associated gene sets. Fisher’s exact test; dotted red line indicates FDR-corrected P<0.05. Cell type/module colors: RG, orange; IPC and Newborn, blue; EN, green; IN, purple. (**B**) Bar and upset plots of Ts21 EN-L6-CT and EN-L6-IT downregulated genes overlapping enriched disease gene sets. (**C**) Rare variant enrichment in differential eRegulon target genes (Fisher’s exact test, FDR-corrected P-value<0.05). CT/IT-associated eRegulons colored in red. Bars colored by cell type as above. Color scale: –log10(FDR P-value). (**D**) Ts21 DAP contributions to heritability across brain traits/diseases (LDSC). Traits include ADHD, ASD, EA, epilepsy, AD, cortical thickness and total surface area. Color: FDR <0.05 (purple) to <0.0001 (yellow).

We found 22 genes downregulated in EN-L6-CT_2 shared across multiple disorders, including ASD (21), syndromic ASD (7), DD (13) and NDD (10). Many (10/22) have established roles in chromatin modification, remodeling and transcriptional regulation (Figure 7B), and four – *TCF20*, *PACS1*, *KMT2C* and *POGZ* – were shared across all 4 sets of risk genes. Notably, TCF20, KMT2C and POGZ regulate transcriptional programs involved in neuronal differentiation and maturation ^175–177^, while PACS1 mediates protein trafficking and affects synaptic density, ion channel activity, and dendrite growth ^178–180^. Similarly, 9/13 downregulated EN-L6-IT genes shared across DD (9) and NDD (11) are chromatin or transcriptional regulators important for neuronal differentiation and maturation, such as KAT6A and EZH2 (Figure 7B). Intriguingly, while most enriched genes were specific to each neuron subtype, two genes associated with both NDD and DD, *GATAD2B* and *SPTBN1*, were downregulated in both EN-L6-CT-2 and EN-L6-IT neurons. SPTBN1 encodes a spectrin cytoskeletal protein implicated in axon growth and synapse formation ^181,182^ and has also been linked to ASD and epilepsy ^182^. GATAD2B is a component of the NuRD chromatin remodeling complex which functions in neuronal differentiation and function, including regulating expression of CT/IT lineage TFs SATB2 and CTIP2 to fine-tune neuronal specification ^183–186^. Reduced GATAD2B levels are consistent with mixed CT/IT identities in Ts21 observed by IHC (Figure 2J-O) and gene expression programs (**Figure S3F,G**), and enrichment of this gene across NDD, DD and DS suggests that impaired excitatory neuron specification may be a shared molecular mechanism underlying ID.

We next investigated if GRNs disrupted in Ts21 overlapped the same genetic signals. We found that 54/90 (60%) differential eRegulons targeted genes associated with at least one of the tested disorders, with all 54 enriched in NDD and 12-31 enriched in other disorders (Figure 7C). The eRegulons shared across ASD, NDD, and DD were primarily active in IPCs, newborn and excitatory neurons and associated with CT/IT specification. Of the 14 differentially active CT/IT eRegulons (Figure 6I), 12 (86%) were enriched in at least one disorder (SOX4, NFIB, TCF4, FOXP2, TBR1, CUX1, BACH1, CUX2, POU3F1, BHLHE22, PKNOX2 and POU6F2; Figure 7C). Altogether, these findings support that molecular and gene-regulatory programs dysregulated in Ts21 are shared across rare-variant genetic signals of ID and ASD and point to altered neuronal specification.

We then asked whether Ts21 DA peaks or altered GRNs were enriched in heritability of neuropsychiatric disorders or other brain traits relevant to DS. We estimated the partitioned heritability of regulatory regions associated with eRegulons differentially active in each cell type (**Supplementary Table S8B**) and Ts21 DA peaks (**Supplementary Table S7A**) by stratified linkage disequilibrium score regression using summary statistics from common variant association for ASD ^187^, ADHD ^188^, Intelligence ^189^, Educational attainment (EA) ^190^, Epilepsy ^191^, Alzheimer’s disease (AD) ^192^, average cortical thickness and total surface area (SA) ^193^ identified by genome-wide association studies (GWAS; Figure 7D). We found a significant contribution of Ts21 DA peaks and eRegulons to heritability of intelligence and educational attainment, suggesting that programs influencing cognition are shared with those that cause ID and localized to most cells of the developing neocortex. Consistent with the reported reduced brain volume in Ts21 and our findings of decreased progenitors, we observed that heritability of total surface area or thickness was also enriched in altered Ts21 eRegulons. Interestingly, the heritability of ASD and ADHD, conditions co-occurring with DS at significant rates was significantly enriched in altered Ts21 eRegulons found in developing EN and IN neurons. In particular, heritability of ASD was specifically localized to EN-L6-IT and EN-L6-CT neurons consistent with rare-variant signals observed in the same cells (Figure 7A**,B**). Finally, we did not observe heritability of AD risk enriched in Ts21-altered chromatin or GRNs in line with previous studies finding accessible chromatin in the developing brain does not overlap with genetic signals of neurodegeneration ^24,25,187^ and demonstrating the specificity of this approach.

## Discussion

Though DS features originate from HSA21 trisomy, the molecular and gene-regulatory mechanisms underlying deficits in brain development and cognitive function had remained poorly understood. Here, we identified specific neurodevelopmental changes in neurogenesis, metabolism, EN specification and cell interactions, and defined gene-regulatory programs controlled directly by HSA21-encoded TFs as well as by secondary effectors driving these cellular changes. Furthermore, we identified disease risk mechanisms, including some localized to deep layer neurons, shared between DS and other neurodevelopmental disorders.

Structural changes in brain volume and gyrification are thought to contribute to the cognitive deficits observed in DS. Previous histological studies have come to varied conclusions as to the cell proportions and/or total number contributing to structural changes ^10^. While studies of adult cortex are confounded by early-onset neurodegeneration, studies using mid-gestation tissue have not agreed on whether neurogenesis is inhibited or whether neurons are dying. We find that during mid-gestation, the period of peak neurogenesis, commitment to the neuronal lineage is accelerated in the Ts21 neocortex, resulting in quicker advancement through the neurogenic timeline and reduced progenitor expansion consistent with proposed models of heterochrony in DS explaining reduced cortical thickness and brain volume ^10^. This is supported by our observation that RG proportions are reduced, that upper layer IT neurons are produced earlier and at greater proportions, and that Ts21 neurons are more transcriptionally mature and connected. These cell type changes are driven by an increase in pro-neurogenic gene expression programs. Some changes can be traced back directly to genes encoded on HSA21, such as the E3 ligase TTC3 which specifically blocks the Akt pathway important for proliferation ^48^, or the metabolic genes *NDUFV5* and *NRIP1*, which likely upset the OXPHOS/glycolysis balance important for radial glial differentiation. Other transcriptional programs are likely secondary effects set in motion earlier in development by Ts21, such as changes in ECM-mediated proliferative signaling and cytoskeletal dynamics promoting cell polarity required for neurogenic asymmetric division of RGs. Accelerated differentiation may not be confined to the neuronal lineage, as we also observed increased pro-angiogenic RG-vascular signaling that could lead to more oxygenation and premature neurogenesis.

We also observed a striking increase in upper layer IT neurons overall and at the expense of deep layer CT neurons at early time points, including increased commitment to the IT lineage indicated by a pseudotime shift from the pool of unspecified neurons to those expressing upper layer markers *NKAIN2* and *SATB2*. While this can be explained in part by acceleration of the neurogenic timeline, we find other genetic factors also contribute to promoting IT identity over that of CT. First, deep layer neurons mis-express several canonical TFs important for establishing and maintaining IT identity such as *SATB2*, *CUX2* and *RORB*, while upper layer neurons conversely mis-express CT transcription factors *BCL11B*, *TLE4* and *ZFPM2*. This mixed CT/IT identity can be observed most clearly during early mid-gestation (GW 13-16), where a significantly larger proportion of Ts21 excitatory neurons are double positive for BCL11B and SATB2. Second, regulation of IT identity can be directly tied to Ts21 through an activator gene-regulatory network enriched in newborn neurons where the HSA21-encoded gene *BACH1* drives expression of IT transcription factors *CUX2* and *BHLHE22*, among others. Notably, *BACH1* expression peaks just before that of *CUX2* in pseudotime, consistent with its potential upstream regulatory function. Finally, CT neuron identity may be broadly affected in Ts21, as gene co-expression analyses show that a reduction in CT expression programs is accompanied by upregulation of those exhibiting mixed corticofugal subtype identities of subplate and subcerebral projection neurons. Together these observations suggest that subtle changes in the relative expression levels of TFs specifying neuron identity, originating from HSA21, may be sufficient to drive large changes.

A subset of individuals with DS are diagnosed with ASD and/or ADHD, suggesting that similar molecular mechanisms may be impacted among neurodevelopmental disorders. Indeed, we find that rare and common variants associated with neurodevelopmental disorders, cognitive function and brain structure are also enriched in Ts21. Rare variants associated with disease risk are localized to altered GRNs active in both deep and upper layer neurons, consistent with the role of ENs in neurodevelopmental disorders. In particular, chromatin remodelers and transcriptional regulators harboring rare loss of function mutations associated with ASD, NDD and DD are dysregulated in Ts21 L6 CT and IT, the most mature neurons at this stage of cortical development. As genetic risk for these disorders has been previously localized to both deep and upper layer neurons, the processes shared with Ts21 are likely to also be found in both subtypes with further maturation. Common variant heritability for outcomes related to cognitive function and brain structure were enriched in regulatory regions altered across progenitors, excitatory neurons and inhibitory neurons, highlighting the importance of precise molecular and genetic control in neurodevelopment. Interestingly, common variants contributing to disease risk shared between Ts21 and ADHD and ASD were enriched in regulatory elements within deep layer neurons. These disorders share, among others, deficits in sensory processing, a process which is strongly influenced by L6 CT projections to sensory thalamic nuclei ^194^. Furthermore, mis-specification of corticofugal neurons that we observed in the Ts21 cortex may broadly impact sensory processing, potentially leading to delayed motor and language development of individuals with DS.

Altogether, our study provides a foundation to interpret Ts21 alterations across model systems and through the developmental timeline to adulthood. Single-nuclei molecular profiling in postnatal ^171^ and adult ^21^ Ts21 neocortex showed compositional changes consistent with those presented here with increased proportions of upper layer IT neurons. Similarly, neuroinflammatory cascades are broadly recognized in the adult DS brain ^112,195^ and detected at early postnatal periods in the companion study ^171^, a parallel of which we begin to observe during mid-gestation in the form of activated microglia transcriptomes. These observations suggest alterations during development are maintained throughout life and could pinpoint opportunities for early intervention. In the future, it will be important to conduct similar studies across other time points and brain regions to fully catalogue the sequence of events and identify the most impactful therapeutic strategies for DS.

## Acknowledgements

We thank members of the de la Torre-Ubieta, Stein, Geschwind, Gandal, Bhattacharyya and Sousa laboratories and Drs. Valerie Arboleda and Chongyuan Luo for helpful discussions and critical reading of the manuscript. Brain tissue was obtained from the UCLA CFAR (5P30 AI028697) and Translational Pathology Core Laboratory (TPCL) and from the NIH Neurobiobank. Illumina sequencing was conducted at the UCLA Eli and Edyth Broad Center of Regenerative Medicine and Stem Cell Research Sequencing Core, multi-ome library quality control was performed at the Technology Center for Genomics & Bioinformatics (TCGB), and imaging was performed at the Intellectual and Developmental Disabilities Research Center (IDDRC) Functional Visualization core, University of California, Los Angeles, Los Angeles, CA. Digital karyotyping was conducted at the IGM Genomics Center, University of California, San Diego, La Jolla, CA.

## Funding

National Institute of Child Health and Human Development grant R21HD108606 (LTU) National Institute of Mental Health grant R01MH124018 (LTU)

UCLA Eli and Edythe Broad Center of Regenerative Medicine and Stem Cell Research Rose Hills Foundation Innovator grant (LTU)

UCLA Eli and Edythe Broad Center of Regenerative Medicine and Stem Cell Research Postdoctoral Training Program (SN)

UCLA Jonsson Comprehensive Cancer Center and Eli and Edythe Broad Center of Regenerative Medicine and Stem Cell Research Ablon Scholars Program (LTU)

NIH Biomedical Big Data Training Program grant T32LM012424 (AW)

## Author Contributions

CV, AW and LTU conceived and designed the study. CV, AW, PS, BS, AS, SN, SY, and LTU collected and processed the tissue specimens. CV generated single-nuclei multi-ome libraries. CV and AW processed the single-cell raw data. PZ, LQ, DHG, MJG and JLS assisted with cell demultiplexing analysis and single-cell analysis pipelines. CV performed demultiplexing, single-cell clustering and annotation, cell composition analysis, differential gene expression analysis, gene co-expression network analysis, and gene-regulatory network analysis. AW and CV performed trajectory inference analysis. AW performed digital karyotypic analysis, cell and neuron communication analyses, and disorder enrichment analysis. PS developed the automated image quantification analysis. CV and PS performed IHC and image quantification. NM and JLS conducted the LDSC analysis. SN and YC supported single-cell analyses. JLS advised on statistical analyses. CV, AW and LTU interpreted the data. CV, AW, PS, NM and LTU wrote the manuscript.

## Data availability

Controlled-access snMultiome data was deposited in the Neuroscience Multi-omic Archive (NeMO) archive (accession number pending).

## Methods

### Tissue acquisition, dissection and single-nuclei isolation

#### Sample acquisition

De-identified human mid-gestation brain tissue was obtained from the UCLA Gene and Cell Therapy Core, the UCLA Translational Pathology Core Laboratory, and the NIH NeuroBioBank repository according to IRB guidelines and to the legal and institutional ethical regulations of the UCLA Office of Human Research Protection. Full informed consent was obtained from all of the parent donors. Excluding the trisomy of HSA21 in Ts21 samples, no known major pathogenic CNVs implicated in neuropsychiatric disorders were found in any donors. Control and Ts21 samples were sectioned to obtain neocortex and flash-frozen in liquid nitrogen (for snMultiome) or fixed in 4% PFA, equilibrated in 30% sucrose and embedded in OCT (for IHC) and stored at - 80℃.

#### Nuclei isolation and snMultiome capture

As manufacturer-provided protocols produced low quality nuclei, we optimized a previously described ^196^ two-step nuclei isolation protocol preserving nuclear integrity and molecular and chromatin signatures. Briefly, frozen neocortical sections (∼10-20mg) were homogenized in RNAase-free conditions in 1-2mL of low-detergent lysis buffer (250mM Sucrose, 25mM KCl, 5mM MgCl2, 10mM Tris-HCL pH7.4, 1% BSA, 0.05% Triton-X, 1mM DTT, 1X EDTA-free protease inhibitor, 0.04U/µL RNase inhibitor) using a 2mL dounce homogenizer on ice. Nuclei were pelleted at 600*g* for 8 minutes at 4℃, washed twice in wash buffer (lysis buffer without Triton-X) and resuspended in 1x PBS supplemented with 1% BSA, 1mM DTT, 1x protease inhibitor and 0.05U/µL RNase inhibitor. Nuclei were filtered through a 40µM tip strainer, and an aliquot was stained with Trypan Blue to confirm nuclei quality and counted on a hemocytometer. Nuclei were further permeabilized for DNA tagmentation by the addition of 5x OMNI buffer (50mM Tris-HCl, 50mM NaCl, 15mM MgCl2, 0.5% Tween-20, 0.5% IGEPAL, 0.05% Digitonin, 5% BSA, 1X PBS, 5mM DTT, 1U/µL RNase Inhibitor, 5X Protease Inhibitor) for 1 minute, pelleted at 600xg for 8 min at 4℃ and washed twice in OMNI wash buffer (10mM Tris-HCL, 10mM NaCl, 3mM MgCl2, 0.1% Tween-20, 1% BSA, 1mM DTT, 1U/µL RNase Inhibitor, 1X Protease Inhibitor). The final nuclei pellet was resuspended in 1x Nuclei Buffer (10X Genomics). An aliquot was stained with Trypan Blue to assess nuclei quality and single nuclei suspension and counted again before pooling and loading onto a 10X multi-ome flowcell. To minimize potential effects of technical batches, nuclei from multiple donor samples were pooled (3-4 samples/pool) into a single suspension, loaded onto the same 10X multi-ome flowcell and processed according to the 10X Genomics Next GEM Single Cell Multiome ATAC kit. Approximately 5,000 nuclei per donor sample were targeted for capture. Libraries were sequenced on a NovaSeq 6000 to a targeted average depth of 25,000 read pairs/nucleus each for RNA and ATAC modalities.

#### Sample genotyping

To confirm sample ploidy and sex, genomic DNA was isolated from flash frozen neocortical samples using the Qiagen DNeasy Blood and Tissue kit (Qiagen, 69504) following the manufacturer’s instructions. Genomic DNA was RNAse-treated (RNAse I, Ambion AM2294) for 30 min at 37℃ followed by heat inactivation at 70℃ for 15 min and subsequently purified using the Genomic DNA Clean & Concentrator kit (Zymo Research, D4064). Single nucleotide polymorphism (SNP) genotyping was performed using the Illumina Infinium CoreExome-24 BeadChip at the UCSD IGM Genomics Center. Genotype calling and calculations for B-allele frequencies (BAF), and Log-R-Ratio (LRR) per sample were performed using Illumina GenomeStudio software (v2.0.5). Each sample was assessed for Ts21 using BAF by the absence of SNV distribution at 0.5 (AB alleles) and presence of SNV distributions at 0.33 (AAB alleles) and 0.66 (ABB alleles). SNV BAFs were plotted along all chromosomes to confirm Ts21 and exclude additional aneuploidies in Ts21 and control samples.

To infer ancestry, donor genotype bedfiles were filtered to remove missing genotypes (geno >0.05), rare variants (maf >0.01), genotypes that deviate from the Hardy Weinberg Equilibrium (hwe >1e^-6^) and genotypes with a high SNP missingness rate (mind >0.01). Ancestry was estimated from donor genotype data using PlinkQC ^197^ (v0.3.4), according to the provided vignette (https://meyer-lab-cshl.github.io/plinkQC/articles/AncestryCheck.html). Non-AT or -GC SNPs were removed from both donor and reference and the bedfile was filtered for variants in linkage disequilibrium with r^2^ > 0.2 in a 50kb window. To ensure genotype bedfiles matched the reference, chromosome and position mismatches were identified and corrected. Allele flips were also identified and either corrected or removed in the absence of a match. Once both were prepared, the bedfile and population reference were merged and principal component analysis (PCA) was run with Plink. The resulting eigenvalues were plotted and analyzed for clustering and association using ggplot2 in R.

### Single-nuclei multi-omic analyses

#### Data pre-processing and sample demultiplexing

CellRanger ARC (v2.02, 10X Genomics) was used to generate fastq files from raw BCL sequencing files for the ATAC modality (cellranger-arc mkfastq), and the cellranger-arc count pipeline was used to process both gene expression and chromatin accessibility fastq files for read mapping, cell barcode calling and generation of quality control metrics according to standard 10X Genomics guidelines using default settings. Reads were aligned to the 10X Genomics pre-built human reference genome (GRCh38 2024-A-2.0.0). SNP-based demultiplexing was used to identify and remove multiplets and assign nuclei to donors. Cellsnp-lite (v1.2.3) ^198^ was used to generate single-nuclei SNP profiles from reads mapped to both RNA and ATAC modalities for each sequencing library. Reference donor SNP profiles imputed from microarray genotyping of exonic variants (Infinium Core Exome-24, see above) were generated using the TOPMed imputation server (https://imputation.biodatacatalyst.nhlbi.nih.gov) and the TOPMed reference panel (v.R3 on GRCh38) ^199^. RNA and ATAC nuclei SNP profiles were matched to reference donor profiles using vireoSNP (v0.5.8) ^200^. Multiplets – nuclei assigned to more than one donor (∼16%) – and those with mismatched donor assignments between modalities (∼1%) were removed. Nuclei with a donor assignment in only one modality (<1%) were retained and assigned using that modality. The majority (∼83%) of nuclei were singlets and assigned to the same donor in both modalities. These were used for all subsequent analyses. Following donor assignment, to minimize effects of cell-free RNA in the medium and ensure accurate transcript quantification, ambient RNA removal was performed on the RNA modality using CellBender (v0.3.0) ^201^, and the resulting nucleus x gene matrix was used for further analyses.

#### Quality control, integration, clustering and cell type annotation

After pre-processing, we applied the following filters to retain high quality nuclei: 1) gene expression count (nCount_RNA) > 200 and < 3 standard deviations from the donor mean, 2) ATAC fragment count (nCount_ATAC) > 100 and 3) percent mitochondrial genes < 5%. The *sctransform* normalization method in Seurat (v4.4.0) ^130^ was used to correct technical variance in read depth and gene detection in individual donor samples. Differences in cell health were corrected for by regressing out mitochondrial gene percentage using var.to.regress in *sctransform*. To control for inter-donor variability and other technical effects, samples were then integrated across donors using the RNA modality by reciprocal PCA projection method with 3000 features (genes). Dimensionality reduction by principal component (PC) analysis was used to minimize technical noise in clustering, and the first 30 PCs were selected for UMAP embedding and clustering. Nuclei were clustered using the Louvain algorithm at a resolution of 0.5. Cluster-enriched genes were identified for each cluster using the *FindAllMarkers* function (min.pct=0.25 and logfc.threshold=0.25) in Seurat. Clusters were annotated by enrichment of cell type markers from previously published datasets of mid-gestation neocortex ^25,26^ (Fisher’s exact test, FDR-corrected P<0.05, with all detected genes in each dataset as background), and by expression of canonical cell type markers. To integrate RNA and ATAC modalities, ATAC read fragments from individual donor samples were first merged, keeping unique cell barcode and donor information, and narrow peaks were called on the merged fragments using MACS2 (v2.2.9.1) with default parameters ^202^. Regions overlapping ENCODE blacklisted regions were excluded. The resulting nucleus x peaks and nucleus x genes matrices were then integrated together using the latent semantic indexing (LSI) method in Signac (v1.12.0) ^203^ to integrate peak and gene embeddings. Nuclei were then re-clustered using the weighted nearest neighbor (WNN) algorithm ^130^ (1-30 PCs and 2-40 LSI components), joint UMAP embedding of the resulting nearest-neighbor graph and Louvain clustering (resolution=1.1). Cell type annotation of resulting clusters was performed as described above for gene expression clusters.

#### Differential cell composition

We employed a generalized mixed model to assess cell composition between conditions. The percent composition of each cell type was first calculated for every donor. For each cell type, generalized mixed models were then fit to the data using the *glmer* function from the lme4 R package ^204^ (v1.1-35.5), with cell type percentage as the output, donor as a random intercept, sex and gestational age as fixed effects, and with or without the condition (control or Ts21) as a predictor. Model fits were compared by ANOVA, and P-values were FDR-adjusted across cell types. For differential composition analyses across pseudotime, P-values were FDR-adjusted across all cell types and pseudotime bins.

#### Differential gene expression analysis

Differentially expressed genes were identified per cell type using a negative binomial gamma mixed model from the NEBULA package ^47^ (v1.5.5). This model properly accounts for the hierarchical structure of the dataset to mitigate pseudoreplication bias that can inflate significance in scRNA-seq. NEBULA was run with condition (control or Ts21) as a predictor, donor as a random effect and sex and gestational age as fixed effects. The number of genes detected in each cell was used as the scaling factor for that cell. Resulting P-values were FDR-corrected across all cells of the respective cluster.

#### Differential program/module expression

Differential expression of CT/IT TFs were calculated for Figure S3, and differential expression of hdWGCNA modules (see below) were calculated for Figs. 4 and S5. The UCell package ^205^ (v2.8.0) was used to calculate the mean expression of sets of genes of interest in each cell type. For comparison of CT/IT programs, cells within the EN lineage were selected and mean expression of gene sets was calculated from RNA counts using the *AddModuleScore_UCell* function. For comparison of hdWGCNA modules, the *ModuleExprScore* wrapper function for UCell from the hdWGCNA package was used to calculate mean expression of the top 25 genes by module eigengene connectivity (kME, see below) for each module. For each cell type, mixed-effects models were fit to the data using the *lmer* function from the lme4 ^204^ R package (v1.1-35.5), with expression values as the output, donor as a random intercept, sex and gestational age as fixed effects, and with or without the condition (control or Ts21) as a predictor. The models were compared by ANOVA, and P-values were FDR-adjusted across cell types.

#### Inferred cell-specific trajectory analysis

Trajectory inference analysis in the EN lineage was performed using Monocle3 ^40,206,207^. The UMAP coordinates and dimensions calculated for the RNA Seurat object (Figure 1A) were used to calculate inter-cell euclidean distances and establish an unordered continuum, or principal graph, with nodes, along which pseudotime was determined. The root was assigned to the node within the highest density of cells that were vRG-oRG-PgS of 13GW donors. Cells that do not differentiate from RGs (i.e. interneurons, microglia, vascular cells) were excluded from pseudotime analysis. Pseudotime was estimated along the trajectory by each cell’s expression profile age relative to the root cells. The significance of the difference in the global trajectory distribution between control and Ts21 cells was determined using the kolmogorov-smirnov test. Within the cell distribution over pseudotime, temporal boundaries or “pseudotime bins” were delimited as the intervals spanning peaks of cell density over pseudotime in control samples. Lineages of EN-Deep and EN-Upper cells were identified by naturally-occurring branches in the principal graph and corresponding similarities in pseudotime estimation. Cell-specific differences in pseudotime were calculated by a pairwise comparison between the estimated marginal means (emmeans) within a linear mixed model (lmer) of pseudotime as a function of condition and celltype, confounding for donor random effects. Results of the pairwise comparison were corrected for multiple testing by FDR.

#### Gene expression over pseudotime

Changes in gene expression for all HSA21-encoded genes and neurodevelopmental TFs were calculated as a function of pseudotime. Normalized expression was extracted for the HSA21-encoded genes and neurodevelopmental TFs per cell, ordered by pseudotime. A smoothing spline was applied to each gene, with 3 degrees of freedom. The resulting gene expression matrix smoothed over pseudotime was normalized to the center of the data by z-score and scaled.

#### Inter-neuronal network estimation and contrast

Inter-neuronal interactions were estimated from the normalized RNA matrix, using NeuronChat (v.1.0.0) ^41^. With the default parameter settings, the inter-neural communication probabilities/strengths and communication network were estimated based on the expression of ligand-receptor pairs and their corresponding weights in the NeuronChat database, per cell type. These ligand-receptor pairs were aggregated into specific neural pathways and their functional similarities were estimated. The resulting individual control and Ts21 NeuronChat networks were contrasted for cell-specific and ligand-receptor specific differential inter-neural communication (FDR corrected <0.05).

#### Cell-network estimation and contrast

Using CellChat (v2.1.2) ^41,208^, the expression of ligand-receptor gene pairs from the CellChat database (CellChatDB v2) was aggregated across the normalized RNA matrix per condition and identified as interactions if sufficiently expressed in sending and receiving cell groups, with default parameters. The expression of ligand-receptor pairs was weighted by network-based smoothing with a PPI adjacency matrix (PPI.human). The interaction strength for each ligand-receptor pair was calculated by trimmed mean summaries of the adjusted expression. The strengths of interactions present in at least 10 cells were aggregated by pathway and then by cell type to generate an interaction network. The centrality, or the contribution, of each cell type to the overall communication network was then calculated. Control and Ts21 CellChat objects were then merged for cross-condition comparison. Within the merged CellChat object, pairwise similarity testing, manifold learning and classification learning were implemented to contrast each condition’s network. The difference in network contribution by pathways was calculated by aggregating pathway signals across cell pairs per condition and assessed by paired Wilcoxon test. Communication probabilities for network pathways were pooled by contributing cell type and transformed by negative reciprocal logarithm (-1/log(p)) to identify cell type-specific differences in network contribution. To assess differences in specific interactions, the merged network was subsetted by ligand-receptor pair and a differential communication probability was calculated by the scaled difference of interaction strengths between conditions by sending and receiving cell type pairs.

#### Gene co-expression network analyses (hdWGCNA)

Single-nucleus high-dimensional weighted gene co-expression network analyses were performed using the hdWGCNA package (v0.3.03) ^72^. hdWGCNA is a systems-level approach which uses unsupervised learning methods to identify groups of co-expressed genes (modules) with differential activity across conditions. It is intrinsically more sensitive to ascertain relevant mechanisms as sub-threshold changes in gene expression can be summed at the module level to reveal dysregulation across conditions. To counter gene drop off inherent in scRNAseq, individual cell type clusters were grouped into 8 major cell types – RG (vRG-tRG, oRG, vRG-oRG-PgS), Pg-Cycling (PgG2M, PgG2M-PgS), IPC (IPC-EN, IPC-EN-Newborn), EN-Newborn (EN-Newborn-1/2, EN-Newborn-IT-Immature), EN-Upper (EN-IT-Immature, EN-L2-L3-L4-IT) EN-Deep (EN-L6-IT, EN-L5-L6, EN-L6-CT_1/2, EN-L6b, EN-L6b-SP), IN-CGE (IN-dLGE-Immature, IN-CGE-Immature) and IN-MGE (IN-MGE-Immature, IN-MGE-SST). Individual co-expression networks were calculated for each major cell type. For each network, the top 2000 most variable genes were selected and retained for network analysis using the variance stabilizing transformation (vst) method in the Seurat *FindVariableFeatures* function. We also required that selected genes were expressed in at least 5% of cells in each major cell type annotation. Soft-power thresholds were tested using the TestSoftPowers function and the lowest threshold to yield a model fit >0.8 was chosen. Metacells were generated using the following parameters: minimum module size=50, maximum cells shared between modules=10, number of nearest neighbors k=25 and RNA-based UMAP coordinates. Signed co-expression networks were then constructed using metacells, with a merge cut height of 0.2 for dynamic tree cutting and default settings for remaining parameters, yielding the following number of modules per network, excluding the grey module, – RG, 7; Pg-Cycling, 6; IPC, 6; EN-Newborn, 6; EN-Upper, 7; EN-Deep, 7; IN-CGE, 4; IN-MGE, 3. Summarized module expression values (ME) were calculated across all cells in the dataset using the *ModuleEigengenes* function and corrected across donors with Harmony ^209^. Eigengene-based connectivity (kME) was calculated for each cell using the *ModuleConnectivity* function. For microglia, analyses were performed using all genes expressed in at least 5% of cells due to the small cluster size. The soft-power threshold was chosen as described above. Metacells were generated using the following parameters: minimum module size=2, maximum shared cells between modules=10, number of nearest neighbors k=2 and RNA-based UMAP coordinates. A signed co-expression network, ME and kME were calculated as described above, yielding 4 modules. Modules in all networks were considered expressed above background if the median module expression score across donors was greater than 0.1 in either condition.

#### Differential chromatin accessibility and motif enrichment analysis

Differentially accessible peaks were identified per WNN cell type via the FindMarkers Seurat function using a logistic regression (LR) model with peak read depth (nCount_peaks), sex and gestational age as latent variables. Peaks with a logFC >0.1 and Bonferroni-corrected P-value <0.05 were considered differentially accessible. Cell type-specific TF motif enrichment within DA peaks was performed using a hypergeometric test via the FindMotifs Signac function with default parameters and the JASPAR 2020 ^210^ motif database. Motifs with a Benjamini-Hochberg corrected P-value <0.05 were considered enriched. The RunChromVar ^211^ wrapper in Signac was used to calculate cell type-specific motif activity scores with default parameters.

#### Enrichment of peaks in epigenetically annotated regions

Enrichment of DA peaks within epigenetically annotated regions was performed using the GREAT method ^131,133^ with imputed chromatin states defined by the Roadmap Epigenomics project ^131^ as previously described ^24^. Briefly, the enrichment of DA peaks within an imputed 25-state chromatin state model derived from prenatal brain tissue (E081 and E082) ^132^ was calculated calculated using the ratio between the (#bases in state AND overlap feature)/(#bases in genome) and the [(#bases overlap feature)/(#bases in genome) X (#bases in state)/(#bases in genome)]. Annotated chromatin state region coordinates were lifted over to hg38. Significance was determined using a binomial test and P-values were FDR corrected.

#### Gene-regulatory network analyses

The SCENIC+ pipeline (v1.0a1) ^136^ was used to build GRNs from paired single-nucleus gene expression and chromatin accessibility multi-omic data. As running the pipeline on all nuclei in the dataset was not possible with current method and memory constraints, we used metacells to reduce the number of cells while preserving data structure and retaining rare cell types. The MetacellsbyGroups function in the hdWGCNA ^72^ package was used to construct metacells within each WNN cell type using the following parameters: minimum size=1, maximum cells shared=2, number of nearest neighbors k=12 and RNA-based UMAP coordinates. The resulting metacell gene expression matrix was used as input into the SCENIC+ pipeline. ATAC fragments were re-assigned to metacells, then MACS2 (v2.2.9.1) was used to call pseudobulked peaks within each cell type. Peaks were extended 250bp on either side and pycisTopic (v2.0a0) was used to derive consensus peaks. High quality metacells were retained using the following filters: minimum unique fragments ≥10^4^ and minimum TSS enrichment ≥7.5. Topic modeling of co-accessible regions (peaks) using the LDA method was then performed for a range of topic numbers and the optimal topic number (30) was chosen as the lowest number where maximum topic coherence and log likelihood converged. Putative enhancer regions were selected using three methods: Otsu thresholding, selecting the top 3,000 regions per topic or identifying differentially accessible regions (Wilcoxon rank sum, logFC >0.5, adjusted p-value <0.05). A custom cistarget database of TF motif scores was generated using Cluster-buster ^212^, the Aerts lab motif collection (v10nr_clust_public) and consensus peaks generated above. This custom cistarget database was used for subsequent motif enrichment within enhancer regions. A SCENIC+ wrapper function was then used to generate eRegulons consisting of a TF, regions enriched for the TF motif, and genes both linked to enriched regions and significantly co-expressed with the TF. High quality eRegulons were retained using the following filters: eRegulons with >10 genes, a positive region-gene relationship and extended eRegulons were kept only when the direct one was not available.

Differentially active eRegulons were identified within each cell type using gene activity. First, area under the curve (AUC) activity scores were calculated for each eRegulon in each metacell and z-score normalized. The mean eRegulon z-score AUC was then calculated per cell type and condition (control or Ts21). An eRegulon with mean z-score>0 in either condition was considered expressed in that cell type. For each expressed eRegulon in each cell type, the fold change was calculated and significance was assessed using a two-sided t-test with equal variance. P-values were FDR corrected within cell type using the number of expressed eRegulons. We required a mean z-scored AUC >0.7 in the control condition and an absolute differential z-score >0.08 to filter out lowly expressed eRegulons and small changes in activity. eRegulons passing these filters and with an FDR-corrected P-value <0.05 were considered differentially active.

### Pathway enrichment analysis

Enrichment analysis was performed using gprofiler2 for Gene Ontology (GO) databases against all expressed genes as the background gene set. For hdWGCNA modules, genes were ranked by highest correlation to the eigengene (kME) and an ordered query was run. For cell type- or class-specific DEGs and eRegulon genes, an unordered query was run. An FDR-adjusted P-value cutoff of 0.05 was used to determine significantly enriched GO terms or pathways and term size was set to <1,000 genes.

### Rare variant enrichment analyses

Gene sets of rare variants associated with neuropsychiatric and neurodevelopmental disorders co-occuring with DS including ASD ^172^, NDD ^172^, DD ^173^, and epilepsy ^174^, as well as those from the SFARI database for syndromic ASD and non-syndromic ASD (score 1) were compiled. Altered Ts21 genes and pathways, including cell-specific DEGs and co-expression modules were tested against each disorder. Cell type and module-specific enrichment with disorders was determined by a Fisher’s exact test, using all expressed genes as the background gene set. To rule out spurious associations, we required each list to have at least 30 genes and at least 5 intersecting genes and an FDR-corrected P-value < 0.05. The same enrichment analysis was applied to significantly differential directed activator eRegulon target genes.

### Partitioned heritability analysis

To evaluate whether differentially-accessible peaks or regions identified in SCENIC+ analysis significantly contribute to heritability in a cell type-specific manner, we performed Stratified Linkage Disequilibrium (LD) Score regression (S-LDSC) ^213,214^ using the baseline-LD model (v1.2) across seven GWAS traits and each cell type separately. GWAS traits used were ASD ^187^, ADHD ^215^, Intelligence ^189^, Epilepsy ^191^, educational attainment ^190^, Alzheimer’s disease ^192^, average cortical thickness and total surface area ^193^. GWAS summary statistics were converted into .sumstats format described as in(https://github.com/bulik/ldsc/wiki/Partitioned-Heritability). Multiple testing correction across cell types was performed using the Benjamini-Hochberg false discovery rate (FDR-BH) method. Significance was defined as FDR < 0.05.

### Immunohistochemistry, imaging and quantification

Immunofluorescent labeling of neocortical cryosections was carried out using standard procedures optimized for fixed tissue. Briefly, 15 μm coronal sections were permeabilized in 1x PBS with 0.2% Triton-X (PT) and blocked with 10% serum. Sections were incubated overnight at 4°C with primary antibodies followed by three PT washes and incubation with fluorophore-conjugated secondary antibodies for 45 min at room temperature. Sections were then counterstained with DAPI, mounted in ProLong Gold antifade reagent and cured overnight at room temperature before imaging. The following primary antibodies were used: SOX2 (Santa Cruz, sc-17320, 1:500), NEUROD2 (Abcam, ab104430, 1:500), SATB2 (Abcam, ab51502, 1:500) and BCL11B (Abcam, ab18465, 1:250). Stained cryosections were imaged on a Leica DMI8 epifluorescence microscope. Tiled images spanning the cortical wall were captured at 20x magnification to include the ventricular/subventricular zones, intermediate zone, subplate, and cortical plate. Resultant multi-channel images were compiled for subsequent cell segmentation and intensity-based cell classification.

Segmentation masks were generated for each nucleus using CellPose v3.0 ^216^. An average of 4,843 and 4,398 nuclei were segmented per image for SOX2/NEUROD2 and BCL11B/SATB2 analyses, respectively. Background signals and large regional pixel intensity differences within images were minimized in ImageJ v2.16.0 using a standardized rolling-ball subtraction algorithm (ball radius=140 pixels), allowing comparable pixel intensity values between nuclei both within and between images. Raw fluorescent intensities were then extracted for each immunolabeled marker. For each channel, marker-positivity of segmented nuclei was assigned through a combination of k-means clustering (SOX2, BCL11B, SATB2) and fixed thresholding (DAPI, NEUROD2). Cortical layer borders were manually defined based on DAPI+ nuclei density and output as normalized distances along the ventricular zone to cortical plate axis for each image. Nuclei were assigned to layers by mapping each nucleus to normalized depth coordinates along the cortical wall. As accurate automatic segmentation of VZ nuclei at the earliest time points (GW13-14) was not possible, we did not include it in the SOX2/NEUROD2 analysis. For each marker set, the percent of nuclei positive for each marker was calculated, and a binomial generalized mixed model was fit to each marker using the *glmer* function from the lme4 R package ^204^ (v1.1-35.5). Marker percentage was used as the output, donor and image as random intercepts, timepoint as a fixed effect, and condition (control or Ts21) as the predictor. Significance was assessed by ANOVA using the carData R package (v3.0-5) and P-values were FDR-adjusted across all markers in the set.

