## Supplemental Materials for "A single cell multi-omic analysis identifies molecular and gene-regulatory mechanisms dysregulated in the developing Down syndrome neocortex"

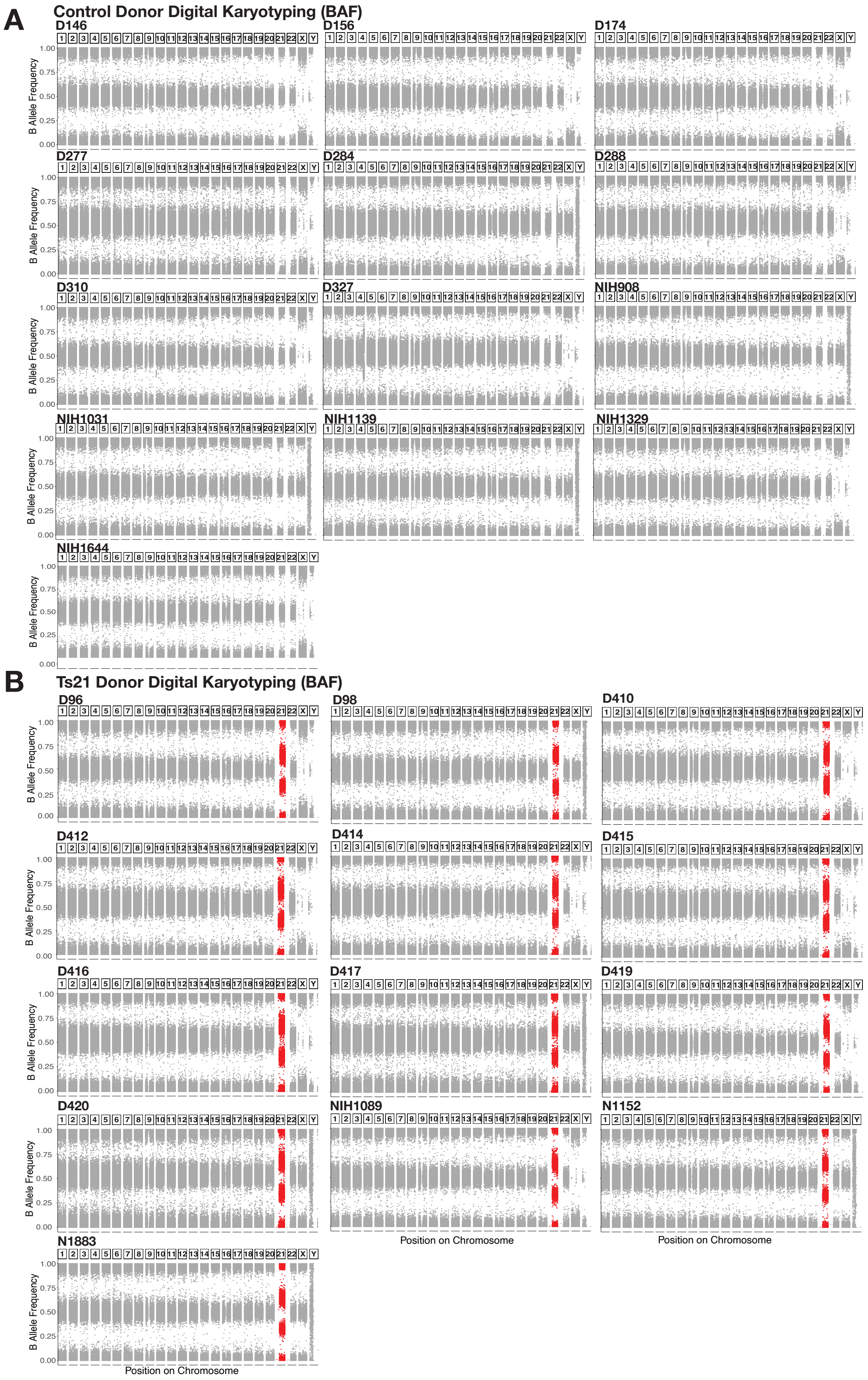


**Figure S1. Digital karyotyping by B-Allele frequency confirms ploidy and sex.**

(**A**) Control pan-chromosomal BAF distributions (D146, D156, D174, D277, D284, D288, D310, D327, NIH908, NIH1031, NIH1139, NIH1329, NIH1644). Chromosomes with BAFs aggregated at 0 (AA), 0.5 (AB), and 1 (BB) convey disomic allele distributions. X chromosomes with BAFs aggregated at 0 (A) and 1 (B) indicate monoallelic distribution, or male. (**B**) Ts21 pan-chromosomal BAF distributions (D96, D98, D410, D412, D414, D415, D416, D417, D419, D420, NIH1089, NIH1152, NIH1883). Non-HSA21 chromosomes with BAFs aggregated at 0 (AA), 0.5 (AB), and 1 (BB) convey disomic allele distributions. HSA21 (red) BAFs aggregated at 0 (AAA), 0.33 (AAB), 0.66 (ABB), and 1 (BBB) convey trisomic allele distributions.

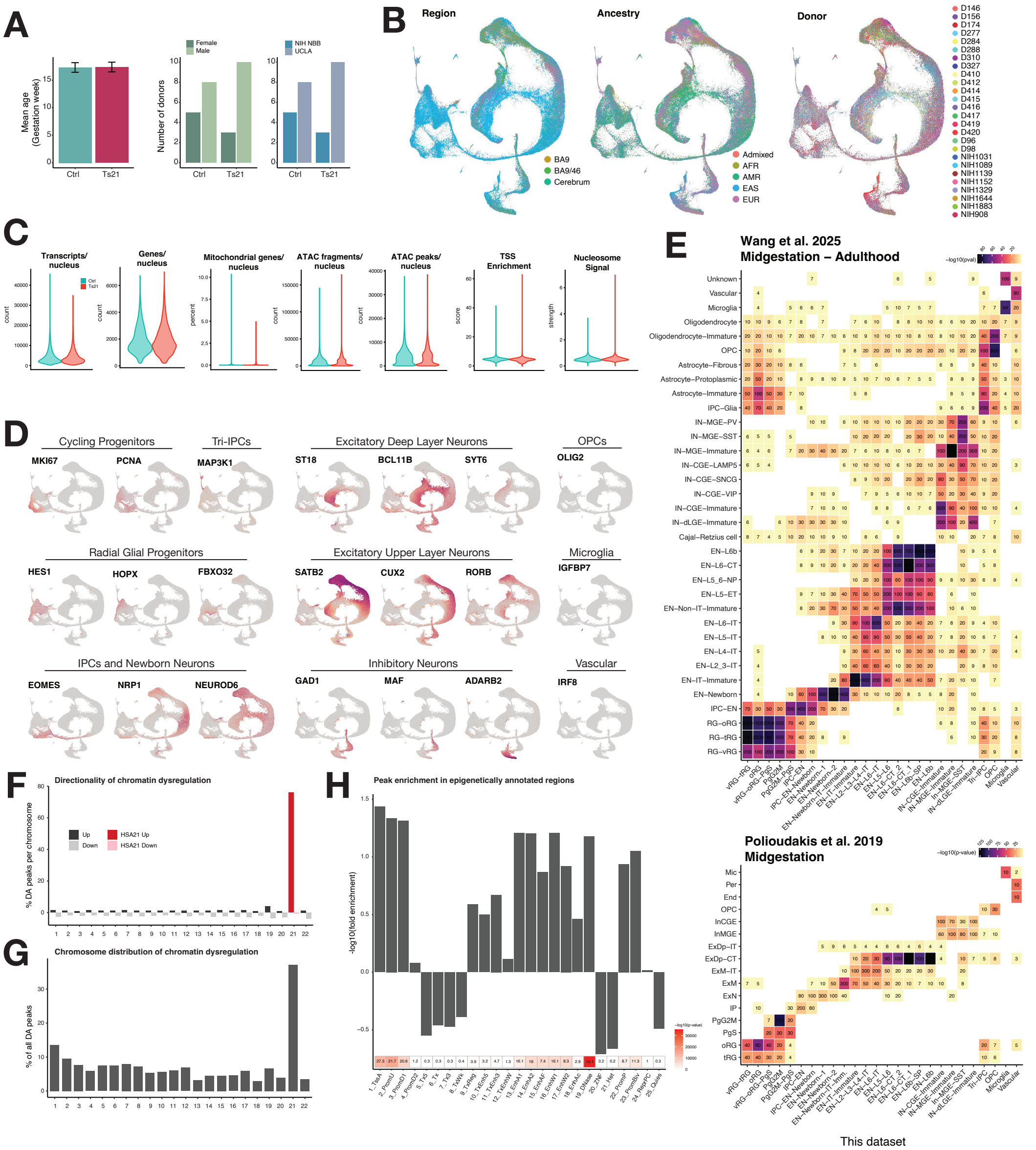


**Figure S2. snMultiome quality control metrics and cell type annotations**.

(**A-B**) Donor cohort metadata. Barplots of mean gestational age, sex and source (A). RNA-based UMAP plots of samples colored by brain region, ancestry and donor (B). Error bars denote standard error of the mean. (**C**) Single-nucleus metadata. Gene expression quality control metrics - per nucleus transcript count, gene count and mitochondrial gene percentage, (left three plots). Chromatin accessibility quality control metrics - per nucleus ATAC fragment count, peak count, transcriptional start site (TSS) enrichment and nucleosome signal strength (right four plots). (**D**) Select cell type-specific marker genes used for cluster annotation. (**E**) Enrichment (Fisher's exact test, FDR-corrected P-value<0.05) of top 50 cell type-specific marker genes from independent human neocortex datasets [25,26] with cluster-specific marker genes in this dataset. Color bars denote -log10 FDR-corrected P-value, numbers indicate odds ratio. **(F)** Differentially accessible (DA) chromatin peaks as a percent of peaks located on each chromosome. Nearly 80% of HSA21 peaks are more open in Ts21 consistent with Ts21. (**G**) DA peaks from each chromosome as a percent of all DA peaks approximately follows chromosome size, except for HSA21. (**H**) Fold enrichment (top) and P-value (bottom) of DA peaks with developing human brain-derived chromatin states shows enrichment in active TSS, promoter and enhancer regions, and under-enrichment in heterochromatin. Color bar denotes -log10 FDR-corrected P-value, number denotes odds ratio.

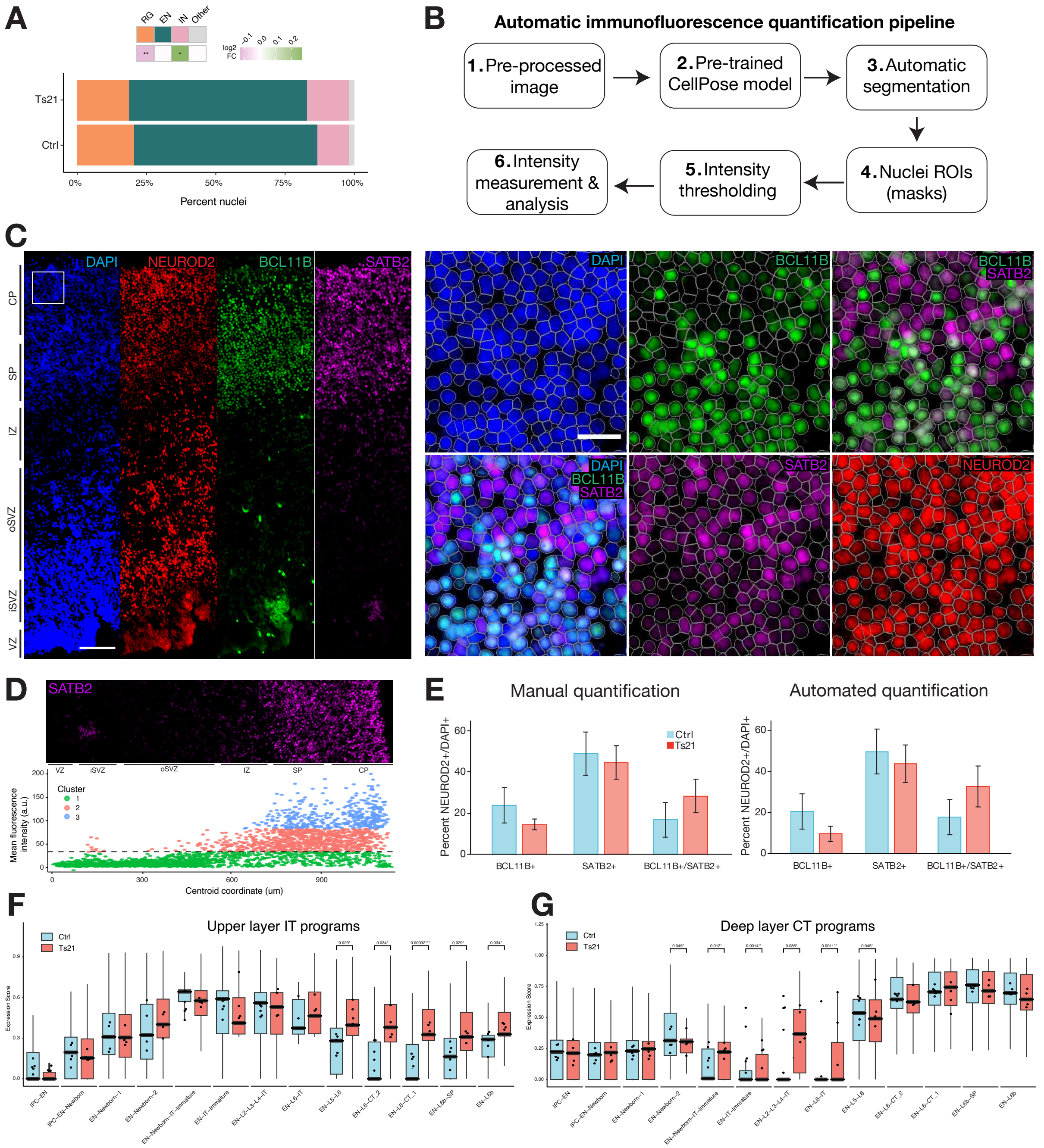


**Figure S3. Semi-automated image quantification pipeline for IHC and IT/CT program mis-expression**.

(**A**) Cell class composition in Ts21 vs control, assessed by linear mixed models accounting for covariates - FDR-corrected *P-value<0.01, **P-value<0.001. (**B**) Automated cell quantification pipeline. (**C**) Representative automatic nuclei segmentation using CellPose. *Left*, cortical strip containing VZ to CP layers. *Right*, inset of CP overlaid with DAPI-based nuclei ROI masks. (**D**) Representative intensity thresholding using k-means clustering. Mean fluorescence intensity of each nuclear ROI is plotted along the length of the VZ to CP axis (x-axis). Dotted black line denotes the chosen minimum fluorescence intensity. (**E**) Comparison of manual versus automated quantification shows nearly identical results. Control, n=3 donors; Ts21, n=4 donors. Error bars denote standard error of the mean. (**F, G**) Mean expression of IT (*SATB2*, *CUX2*, *RORB*) or CT (*BCL11B*, *TLE4*, *ZFPM2*) transcription factors across excitatory neurons. The UCell method was used to calculate average IT or CT program expression. IT programs are significantly increased in deep layer neurons, while conversely CT programs are significantly increased in newborn and upper layer neurons. Linear mixed models, FDR-corrected *P<0.05, **P<0.01, ***P<0.001. VZ, ventricular zone; iSVZ, inner subventricular zone; oSVZ, outer subventricular zone; IZ, intermediate zone; SP, subplate; CP, cortical plate.

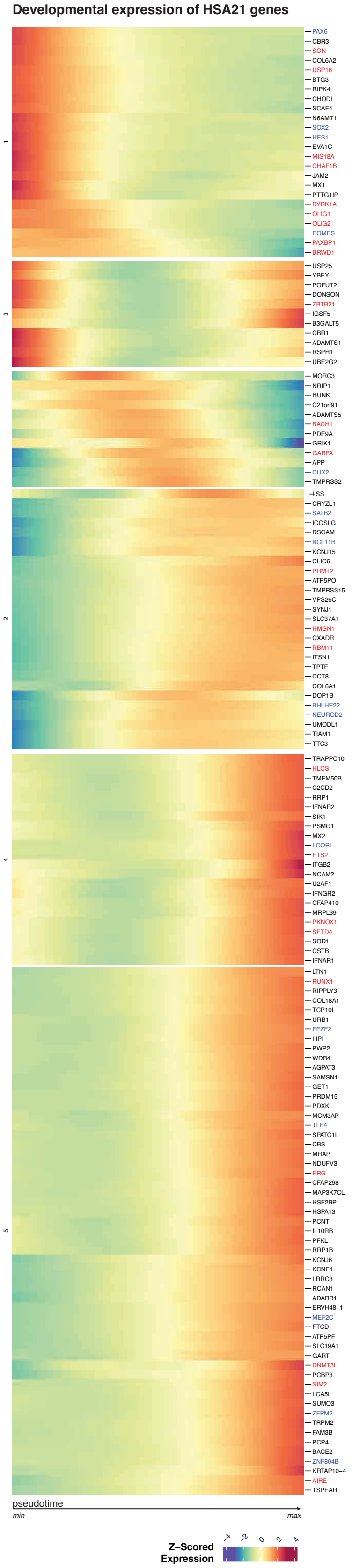


**Figure S4. Expression of HSA21 genes across pseudotime during corticogenesis.**

Normalized expression of HSA21 and select TFs involved in corticogenesis detected in the excitatory neuron lineage over pseudotime. Color bar denotes z-score normalized expression. Select non-HSA21 corticogenesis TFs are labeled in blue, HSA21 TFs or epigenetic regulators are labeled in red. All other HSA21 genes are labeled in black.

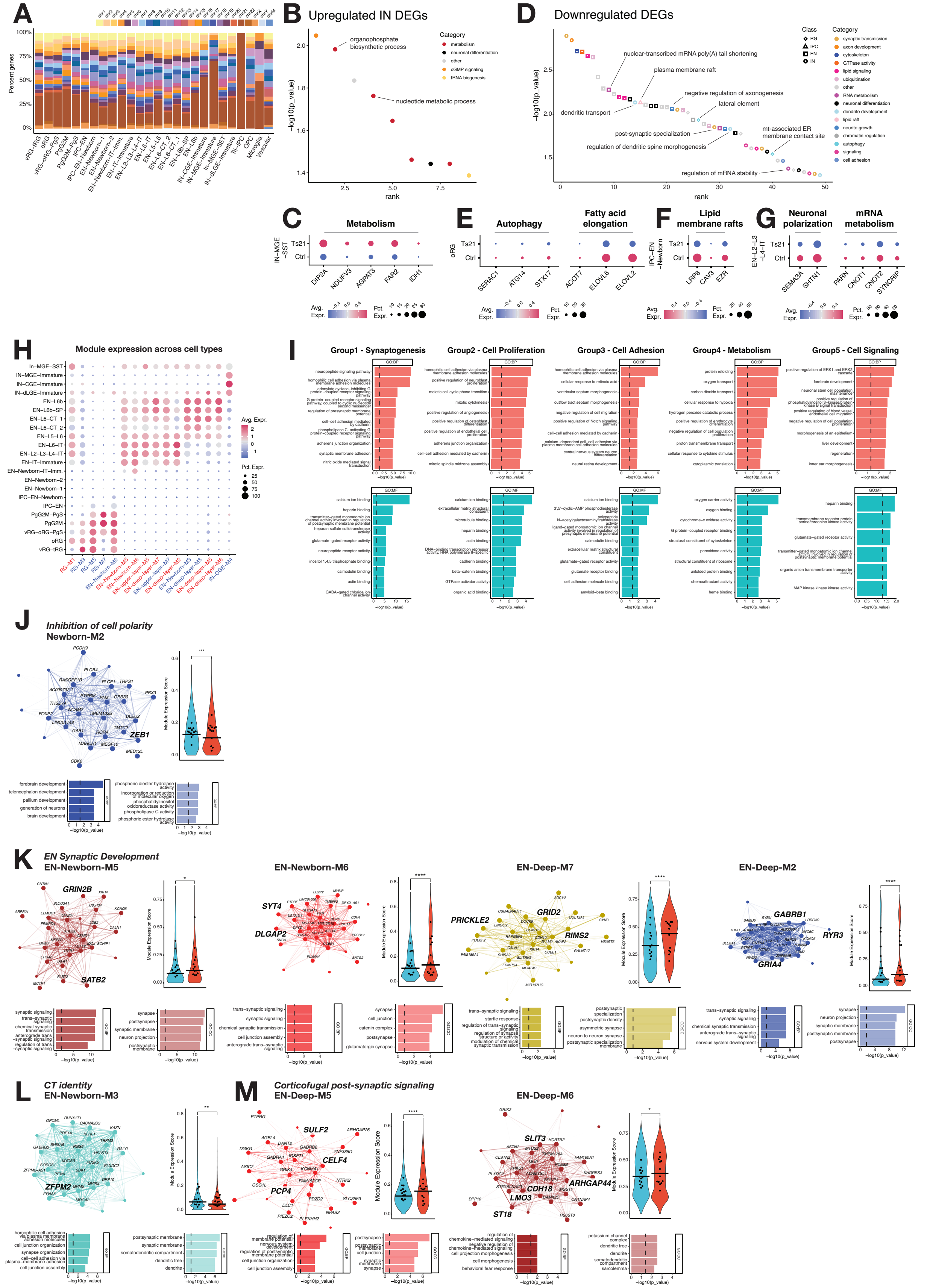

**Figure S5. Differential gene expression and differential co-expression networks.**

(**A**) Chromosomal distribution of cell type-specific differentially expressed genes (DEGs). HSA21 genes comprise ~25% of DEGs per cell type. (**B**) Upregulated IN DEG GO terms ranked by p-value. **C**) Select upregulated genes related to metabolism in IN-MGE-SST. (**D**) Top downregulated GO terms ranked by p-value. Terms are colored by category, with specific GO terms highlighted. Shape indicates cell class (D). (**E-G**) Select genes related to processes enriched in downregulated genes in RGs (E), IPCs (F) and ENs (G). (**H**) Module eigengene expression across cell type-specific clusters. Modules up- and downregulated in Ts21 are indicated in red and blue, respectively. Color bar indicates scaled expression of each module across clusters. Dot size denotes the percentage of cells in each cluster expressing the module eigengene. (**I**) Top 10 gene ontology terms of co-expression module groups. BP, biological process; CC, cell compartment. (**J-M**) Highlighted differential co-expression modules. (**J**) Module Newborn-M2 regulating cell polarity is downregulated in Ts21. **K**) Multiple modules regulating development and organization of pre- and post-synaptic compartments (EN-Newborn-M5/M6, EN-Deep-M7/M2) are upregulated. (**L,M**) Deep layer neuron identity modules. CT identity (L, EN-Newborn-M3) is decreased while subplate (M, EN-Deep-M6) and non-CT identities are upregulated (M, EN-Deep-M5).

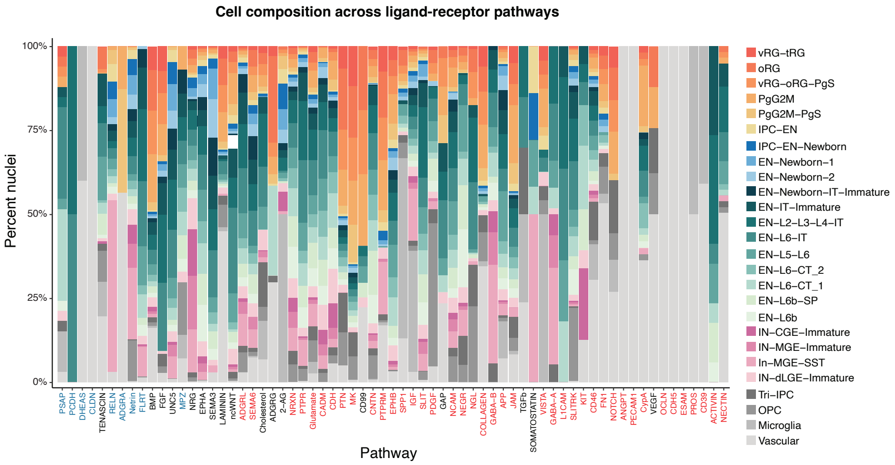


**Figure S6. Cell type contribution to cell-cell interaction pathways.**

Percent nuclei composition of all 67 pathways in the merged network. Pathways enriched in control are labeled in blue, while those enriched in Ts21 are labeled in red. Cell types are indicated by color legend at right.

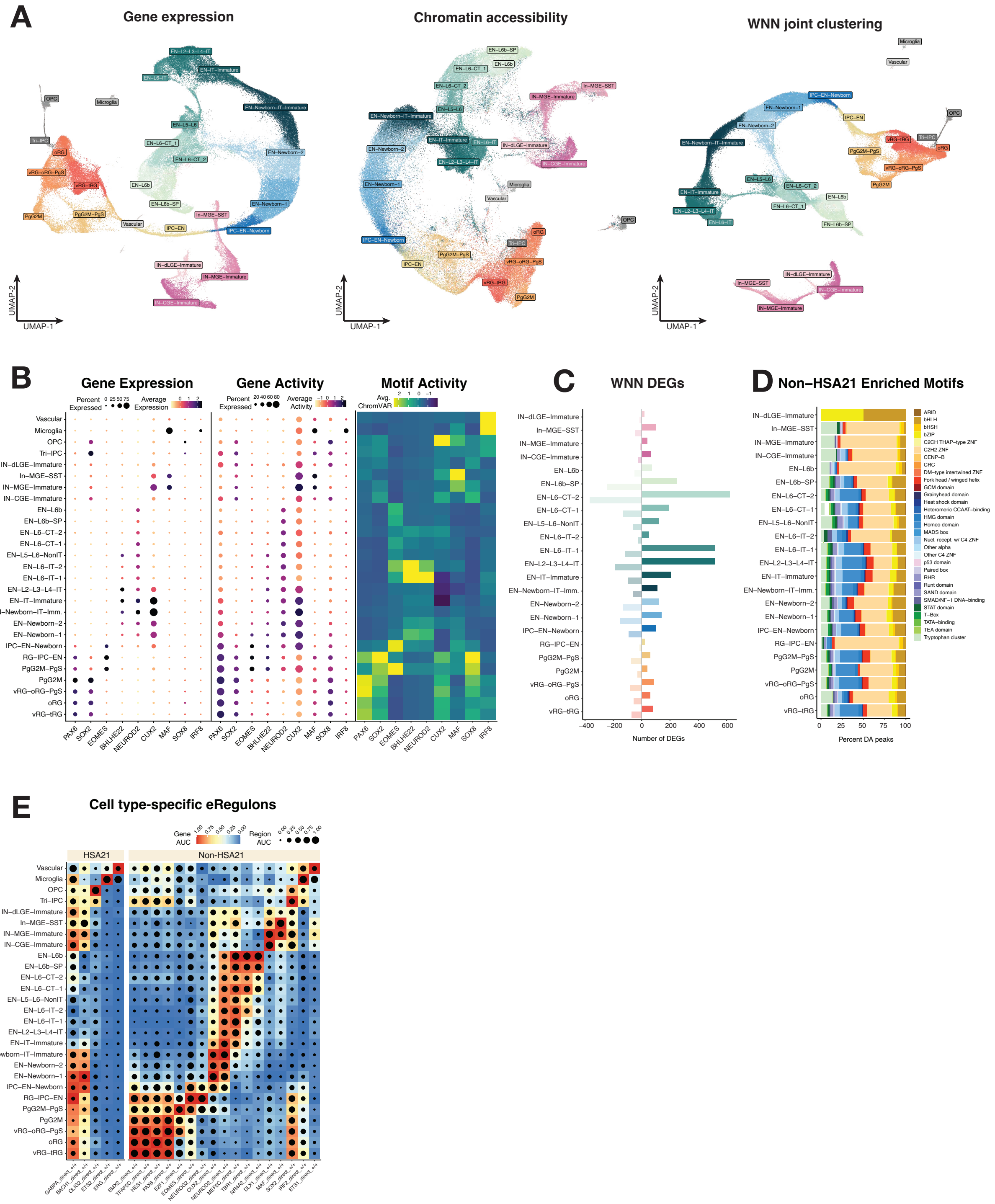


### Figure S7. Concordance between gene expression and chromatin accessibility and GRN fidelity.

(**A**) RNA-, ATAC- and WNN-based UMAPs showing clustering by the respective single or joint modalities. Nuclei are colored by RNA-based annotation. (**B**) Left, expression of select cell type-specific TFs active during neurogenesis. Center, gene activity, or chromatin accessibility at the TSS. Right, normalized motif activity calculated by chromVar. Color bars indicate mean expression or activity, dot size indicates percent of cells expressing the gene or gene-associated accessible chromatin. (**C**) Number of DEGs per WNN cell type. **(D**) Percent of DA peaks enriched for known TF motifs across WNN cell types. TF family is indicated in legend at right. (**E**) Heatmap dotplot of detected HSA21 and select neurogenesis-related TF eRegulon activity (AUC, area under the curve) by target gene expression (color bar) and chromatin accessibility (dot size).

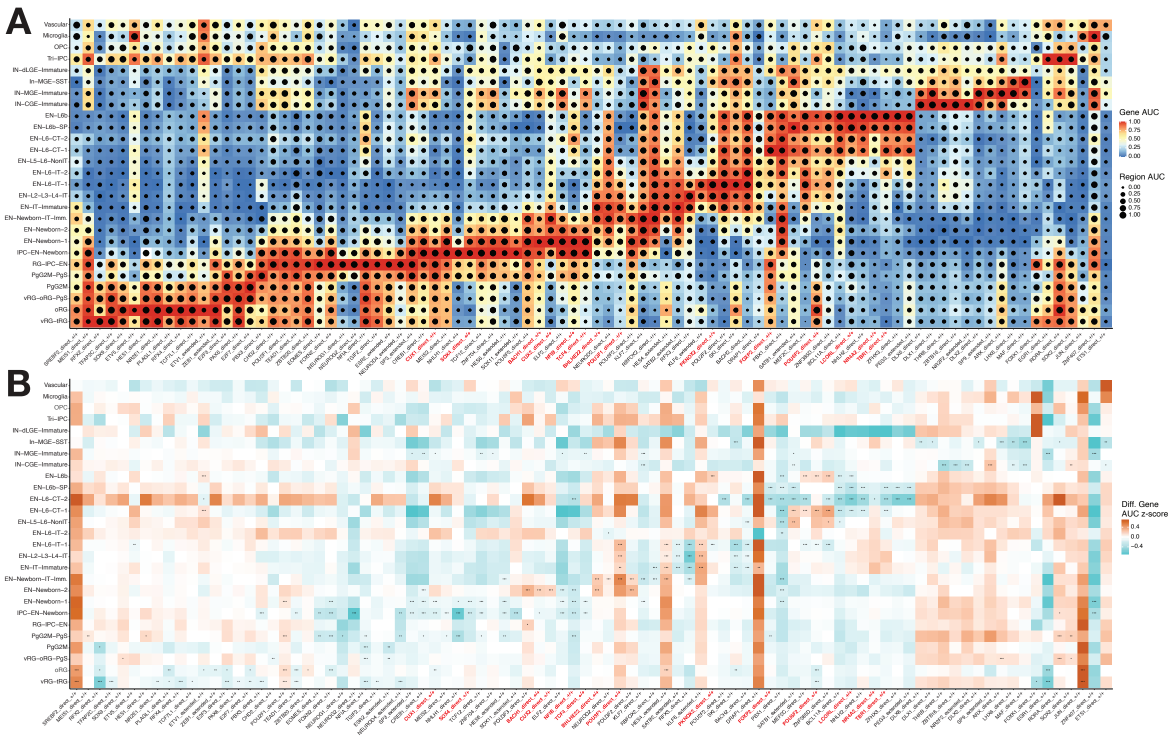


**Figure S8. Cell type-specific differential eRegulons.**

(**A**) Heatmap dotplot of eRegulon activity (AUC, area under the curve) by target gene expression (color bar) and chromatin accessibility (dot size). (**B**) Heatmap of differential eRegulon gene activity. Color bar denotes differential activity z-score. Two-sided t-test, FDR-corrected *P<0.05, **P<0.01, ***P<0.001. CT- or IT-promoting eRegulons are highlighted in red.

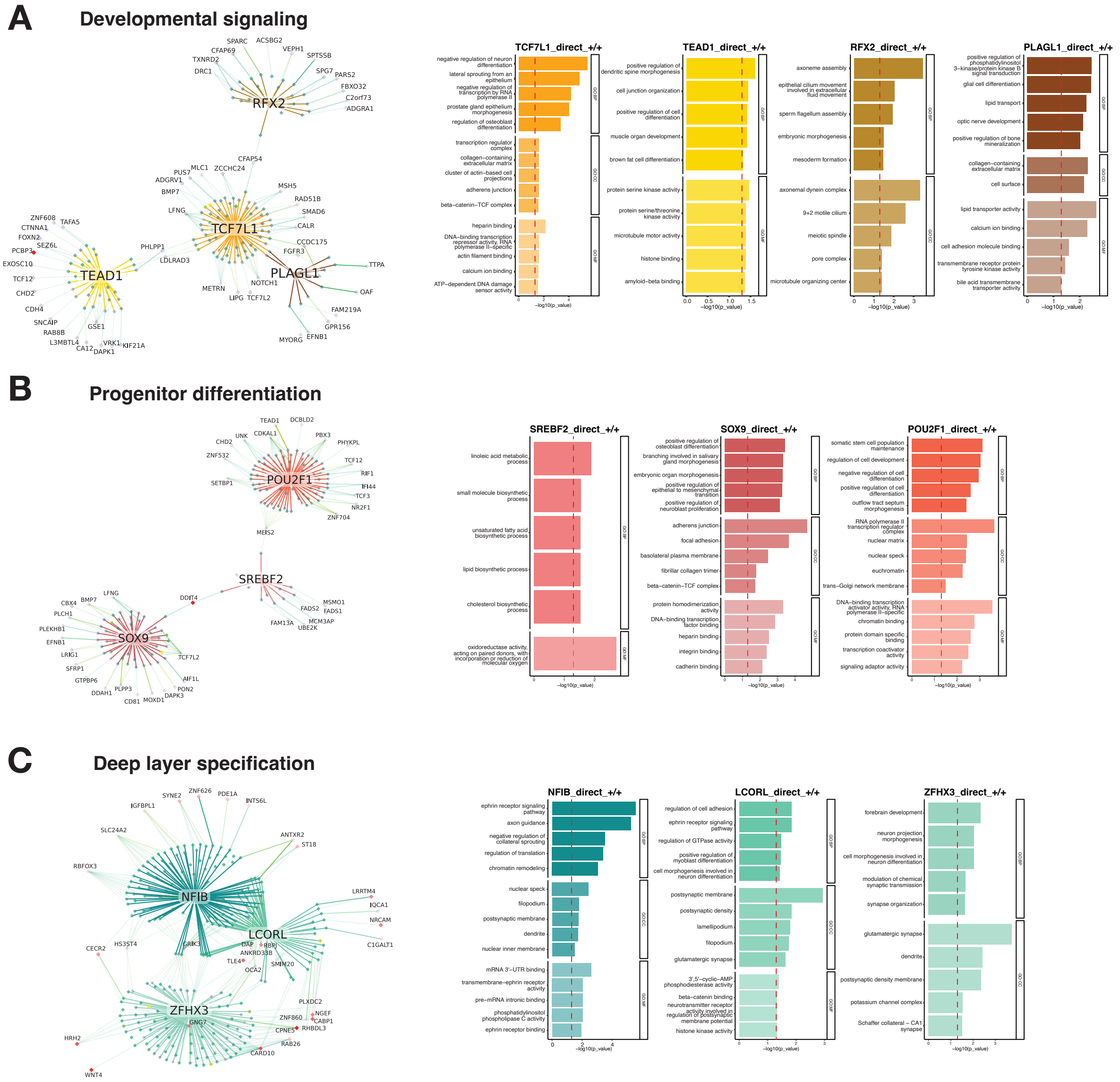


**Figure S9. Additional Ts21 dysregulated GRNs directing neural progenitor signaling and differentiation and deep layer specification.**

(**A**) Left, TCF7L1, TEAD1, PLAGL1 and RFX2 pro-proliferative signaling eRegulons. Primary edges from TFs denote significant TF-to-region connections and inner green diamonds denote chromatin regions. Secondary edges from regions denote significant region-to-gene (R2G) connections, with darker colors and thicker edges indicating a stronger R2G score. Outer diamonds denote eRegulon genes targeted by each region; red color denotes increased gene expression in Ts21. Right, top 5 biological process (BP), cell compartment (CC) and molecular function (MF) gene ontology terms enriched in selected eRegulon target genes. Red dotted line denotes FDR-corrected P-value<0.05. (**B**) Select eRegulons promoting RG differentiation - POU2F1, SREB2 and SOX9. (**C**) Select eRegulons involved in CT identity (NFIB and LCORL) or ID (ZFHX3).

**Supplementary Table S1. Single nucleus metadata**

(A) Donor metadata; (B) Single nucleus metadata.

**Supplementary Table S2. Differential gene expression (NEBULA)**

(A) Cell type-specific NEBULA results, fdr<0.1.

**Supplementary Table S3. Pseudotime analysis**

(A) Single cell pseudotime values and metadata.

**Supplementary Table S4. Neuron communication analysis (NeuronChat)**

(A) NeuronChat control Network Communication Probability; (B) NeuronChat Ts21 Network Communication Probability; (C) NeuronChat Network Differential Communication Probability; (D) NeuronChat Information Flow.

**Supplementary Table S5. Gene co-expression networks (hdWGCNA) and differential analyses**

(A) RG Network; (B) Pg-Cycling Network; (C) IPC Network; (D) EN-Newborn Network; (E) EN-Deep Network; (F) EN-Upper Network; (G) IN-CGE Network; (H) IN-MGE Network; (I) Differential Modules – RG, Pg-Cycling, IPC, EN, IN; (J) Microglia Network; (K) Differential Modules – Microglia.

**Supplementary Table S6. Cell-cell interaction analysis (CellChat)**

(A) CellChat Information Flow (RankNet); (B) CellChat IntraVascular Interactions Differential Communication Probability; (C) CellChat Vascular Interactions Differential Communication Probability; (D) CellChat Microglia Interactions Differential Communication Probability.

**Supplementary Table S7. Differential chromatin accessibility analyses**

(A) Cell type-specific differentially accessible peaks; (B) Motif enrichment in DA peaks.

**Supplementary Table S8. Gene-regulatory networks (SCENIC+) and differential analyses**

(A) Detected eRegulons; (B) Cell type-specific differential eRegulons.

**Supplementary Table S9. Genetic enrichments**

(A) Rare variant enrichment in DEGs and co-expression modules; (B) Rare variant enrichment in eRegulons; (C) Common variant enrichment (LDSC) in differential eRegulons and DA peaks.
